# Enhanced fungal specificity and *in vivo* therapeutic efficacy of a C-22 modified FK520 analog against *C. neoformans*

**DOI:** 10.1101/2023.06.05.543712

**Authors:** Angela Rivera, Won Young Lim, Eunchong Park, Patrick A. Dome, Michael J. Hoy, Ivan Spasojevic, Sheng Sun, Anna Floyd Averette, Sergio Pina-Oviedo, Praveen R. Juvvadi, William J. Steinbach, Maria Ciofani, Jiyong Hong, Joseph Heitman

## Abstract

Fungal infections are of mounting global concern, and the current limited treatment arsenal poses challenges when treating such infections. In particular, infections by *Cryptococcus neoformans* are associated with high mortality, emphasizing the need for novel therapeutic options. Calcineurin is a protein phosphatase that mediates fungal stress responses, and calcineurin inhibition by the natural product FK506 blocks *C. neoformans* growth at 37°C. Calcineurin is also required for pathogenesis. However, because calcineurin is conserved in humans, and inhibition with FK506 results in immunosuppression, the use of FK506 as an anti-infective agent is precluded. We previously elucidated the structures of multiple fungal calcineurin-FK506-FKBP12 complexes and implicated the C-22 position on FK506 as a key point for differential modification of ligand inhibition of the mammalian versus fungal target proteins. Through *in vitro* antifungal and immunosuppressive testing of FK520 (a natural analog of FK506) derivatives, we identified JH-FK-08 as a lead candidate for further antifungal development. JH-FK-08 exhibited significantly reduced immunosuppressive activity and both reduced fungal burden and prolonged survival of infected animals. JH-FK-08 exhibited additive activity in combination with fluconazole *in vivo*. These findings further advance calcineurin inhibition as an antifungal therapeutic approach.

**Importance:** Fungal infections cause significant morbidity and mortality globally. The therapeutic armamentarium against these infections is limited and development of antifungal drugs has been hindered by the evolutionary conservation between fungi and the human host. With rising resistance to the current antifungal arsenal and an increasing at-risk population, there is an urgent need for the development of new antifungal compounds. The FK520 analogs described in this study display potent antifungal activity as a novel class of antifungals centered on modifying an existing orally-active FDA approved therapy. This research advances the development of much needed newer antifungal treatment options with novel mechanisms of action.

## Introduction

Fungal infections, which result in more than 1.5 million deaths per year, are recognized as a serious global health threat (1). The World Health Organization recently released a prioritized list of fungal pathogens for global research (2). The critical priority group consists of 4 life-threatening fungal pathogens: *Cryptococcus neoformans, Candida albicans, Candida auris,* and *Aspergillus fumigatus* (2). Underscoring the global severity of *C. neoformans* infection, cryptococcal meningoencephalitis, is responsible for ∼20% of all AIDS-related deaths, and is among the deadliest fungal infections of global impact (3). These infections have a further devastating impact on sub-Saharan African populations (1, 3, 4). The current treatment for cryptococcal infection consists of combination therapy with amphotericin B and flucytosine in high-income countries; however, those most impacted by this health burden have fewer and less effective treatment options (5). Flucytosine is typically not available in these regions, and often fluconazole alone is prescribed due to its oral availability. However, fluconazole antifungal activity against *C. neoformans* is only fungistatic, and resistance frequently develops in the treated patient as a result of aneuploidy (6). Resistance to fluconazole is rapidly emerging in multiple other fungal species including *Candida auris* and *Aspergillus fumigatus* that is in part due to environmental use of azoles in agriculture, further limiting this therapeutic option and driving the need for newer antifungal drugs (7–9).

One promising antifungal drug target under investigation is the protein phosphatase calcineurin. Calcineurin is highly conserved across pathogenic organisms including fungi and parasites (10–12). Calcineurin mediates stress responses in *C. neoformans, C. albicans,* and *A. fumigatus* (13–16). Calcineurin inhibition by FK506 prevents *C. neoformans* growth at high temperature (17, 18). In fact, calcineurin is essential for virulence in *C. neoformans* and also in several other pathogenic fungal species including *Candida* and *Aspergillus*, and therefore is an attractive antifungal drug target (15, 19, 20). The high degree of conservation of calcineurin across human fungal pathogens supports the approach of targeting calcineurin to develop a pan-antifungal therapeutic.

However, calcineurin is also conserved in humans (21). Inhibiting calcineurin in humans results in immunosuppression. Specifically, the natural calcineurin inhibitor FK506 (also known as tacrolimus) induces immune suppression by inhibiting T cell activation and interleukin-2 (IL-2)-dependent responses mediated by the nuclear factor of activated T cells (NFAT) transcription factor (22–28). Therefore, in this study we monitored IL-2 expression as a measure of mammalian calcineurin inhibition. The FK506 ligand first binds to the FK506-binding protein (FKBP12) and the FKBP12-FK506 complex then binds to and inhibits calcineurin (29–31). FK506 is an FDA approved therapeutic initially deployed to prevent graft rejection following organ transplants (32, 33). FK506 has also been researched for use in additional applications including during treatment of hepatic fibrosis, glioblastoma, as well as aging and bone degeneration in Alzheimer’s (34–37). Currently, FK506 has received additional FDA approval for use in atopic dermatology applications (38).

Humans and fungi share a high degree of conservation of cellular machinery making it difficult to identify fungal-specific drug targets. As such, immunosuppression by FK506 precludes its use as an antifungal therapeutic as immune suppression is a factor that increases the risk of fungal infection (39). Our goal is to modify the structure of the calcineurin inhibitor FK520 (a natural product analog of FK506 or ascomycin) to preserve antifungal activity and reduce or eliminate immunosuppressive activity. Both FK506 and FK520 inhibit *C. neoformans* growth under stress conditions, and synthesis of less or non-immunosuppressive analogs that retain antifungal activity is a validated antifungal drug development approach (40–43). We previously solved high resolution X-ray crystal structures of the calcineurin-FK506-FKBP12 complexes from fungal pathogens including *A. fumigatus, C. albicans,* and *C. neoformans.* Using structure-guided drug design, the carbon at position 22 (C-22) on FK506 was implicated as a critical point for chemical modification to achieve differential ligand targeting of human and fungal calcineurin (44). Thus far, it was demonstrated how modifications at this site on both FK506 and its natural analog FK520, differing by a single allyl to ethyl group substitution at the C-21 position, resulted in decreased immune suppression while maintaining sufficient antifungal activity to be of benefit in animal infection models (45, 46). Developing FK506 analogs as an antifungal therapeutic approach is further bolstered by the research describing synergistic activity of calcineurin inhibitors with additional antifungal agents currently being investigated in clinical trials (47).

Here we have developed an expanded library of FK506/FK520 analogs modified at the C-22 position. These efforts have resulted in the generation of compound JH-FK-08 as a novel antifungal therapeutic effective at significantly reducing lung fungal burden in a murine cryptococcosis model compared to untreated infected animals. Furthermore, treatment with JH-FK-08 significantly prolonged murine survival and surpassed the antifungal efficacy achieved in previous efforts. These advances identify an FK520 analog with potential to be utilized to treat invasive fungal infections.

## Results

### Chemical synthesis of novel FK520 analogs modified at the C-22 position

Previous studies determined the C-22 position on FK506/FK520 as a key point of differentiation between fungal versus human calcineurin-FK506-FKBP12 complexes (44, 45). To selectively modify the C-22 position on FK520, we explored the condensation reaction with hydrazine derivatives. The condensation reaction with hydrazine derivatives exploits the unique reactivity of carbonyl groups, such as ketones and aldehydes, towards hydrazine derivatives in the presence of other functional groups. Based on this rationale, 12 new FK520 analogs were synthesized through one-step syntheses that can be readily scaled up for studies in animal infection models for pre-clinical analysis (Fig. 1) (44, 48). The synthesis of FK520 analogs with C-22 modifications features a condensation of FK520 with a hydrazide or a substituted hydrazine either under reflux or with acidic conditions. After completion of the reaction, the reaction mixture was concentrated *in vacuo* and purified by silica gel column chromatography to give the desired C-22-modified FK520 analogs (33-52% yield). Each analog was validated through ^1^H nuclear magnetic resonance (NMR) and high-resolution mass spectrometry (HRMS) (Text S1, Table S1).

**Fig. 1.**
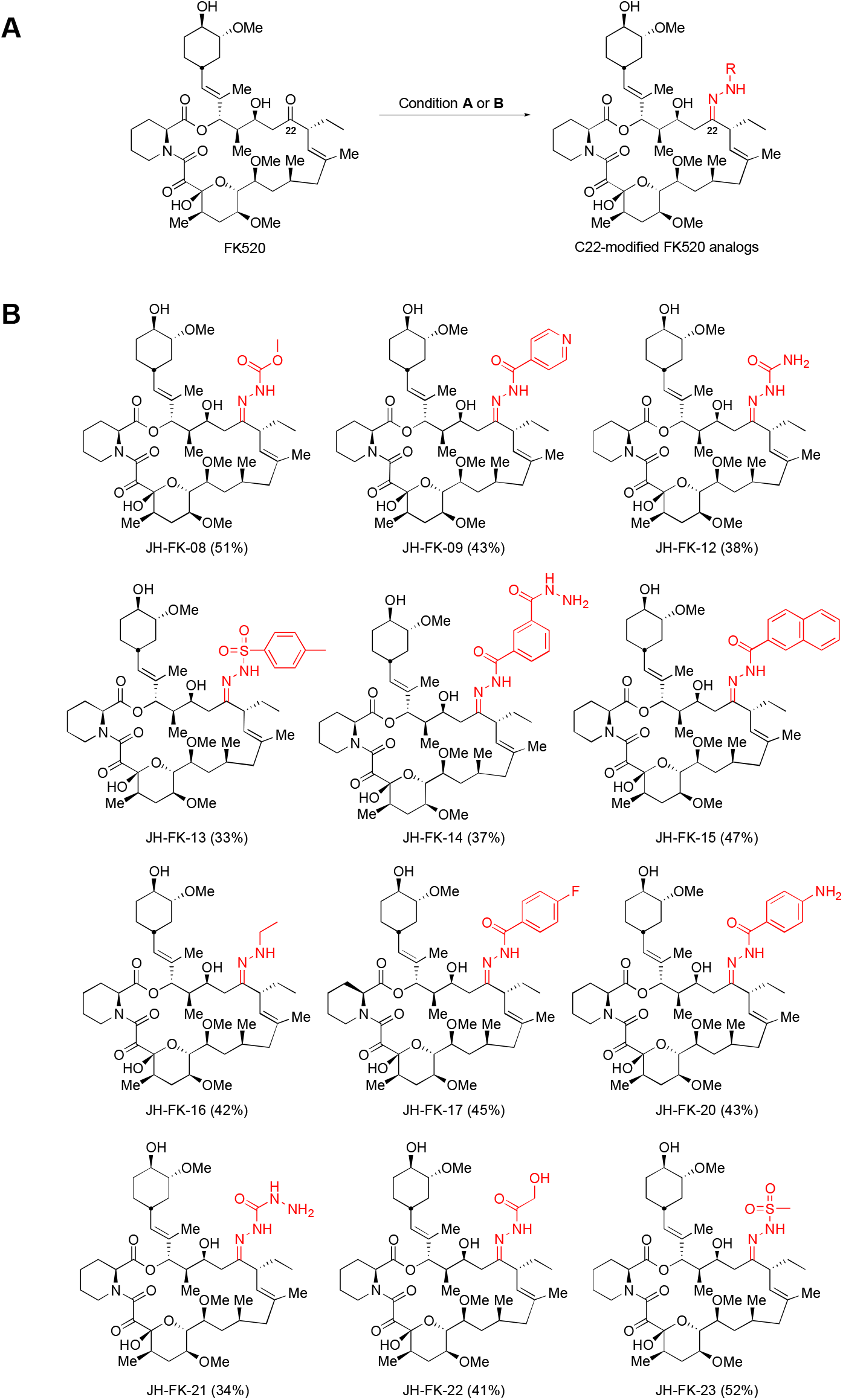
Chemical structures of novel C-22 modified FK520 analogs synthesized in this study. (*A*) One-step synthesis for C-22 modified FK520 analogs. Condition **A**: RNHNH_2_, EtOH, 90 °C, 48 h; Condition **B**: RNHNH_2_, MeOH, TFA (cat), 25 °C, 48 h. (*B*) Structures of 12 analogs modified with a hydrazide or a substituted hydrazine at the C-22 position of FK520 are depicted.

### FK520 analogs display conserved antifungal activity

Eleven of these FK520 analogs were found to retain some level of antifungal activity. To assess antifungal activity, the panel of compounds was tested for fungal growth inhibition using minimum inhibitory concentration (MIC) assays against *C. neoformans* and *C. albicans,* and minimum effective concentration (MEC) assays against *A. fumigatus*. The results from these broth microdilution assays showed a decrease in antifungal activity of the modified compounds compared to FK506 and FK520 (Table 1). Ten of these analogs exhibited an MIC value between 0.4 to 1.6 μg/mL which is an 8- to 32-fold decrease in activity compared to FK506 or FK520 in *C. neoformans* and *C. albicans.* MEC values against *A. fumigatus* ranged from 0.6 to 2.5 μg/mL for 10 of the analogs tested. Notably, analogs JH-FK-13, JH-FK-15, and JH-FK-17 exhibited solubility issues during microdilution broth assays (Fig. S1) (49–51). Overall, a majority of these analogs maintain relatively potent antifungal activity across species and represent an active library for further investigation.

**Table 1.** MICs for FK506, FK520, and analogs*^a^*

| **Compound** | **MIC/ MEC (µg/mL) against:** |  |  |
| --- | --- | --- | --- |
|  | ***Cryptococcus neoformans* (H99)** | ***Candida albicans* (SC5314)** | ***Aspergillus fumigatus* (*akuB*<sup>Ku80</sup>)** |
| FK506 | 0.05 | 0.02 | 0.04 |
| FK520 | 0.05 | 0.02 | 0.16 |
| JH-FK-08 | 0.8 | 0.4 | 1.3 |
| JH-FK-09 | 0.8 | 0.8 | 1.3 |
| JH-FK-12 | 0.8 | 0.8 | 2.5 |
| JH-FK-13 | >50 | >50 | >10 |
| JH-FK-14 | 1.6 | 0.8 | 0.6 |
| JH-FK-15 | 3.1 | 0.4 | 0.3 |
| JH-FK-16 | 0.4 | 0.4 | 1.3 |
| JH-FK-17 | 0.4 | 0.4 | 1.3 |
| JH-FK-20 | 0.8 | 0.8 | 0.6 |
| JH-FK-21 | 1.6 | 0.8 | 2.5 |
| JH-FK-22 | 1.6 | 1.6 | 2.5 |
| JH-FK-23 | 1.6 | 1.6 | 5 |

<sup>a</sup> MIC<sub>90</sub> were tested at 37°C for *C. neoformans*. MIC<sub>80</sub> were tested at 30°C with 2 µg/mL supplementary fluconazole for *C. albicans*. MEC was collected for *A. fumigatus* strains.

**Table 1** MICs for FK506, FK520, and analogs<sup>a</sup>
| **Compound** | **MIC/ MEC (µg/mL) against:**<br>***Cryptococcus***<br>***neoformans* (H99)** | ***Candida albicans***<br>**(SC5314)** | ***Aspergillus fumigatus***<br>**(*akuB*<sup>Ku80</sup>)** |
| --- | --- | --- | --- |
| FK506 | 0.05 | 0.02 | 0.04 |
| FK520 | 0.05 | 0.02 | 0.16 |
| JH-FK-08 | 0.8 | 0.4 | 1.3 |
| JH-FK-09 | 0.8 | 0.8 | 1.3 |
| JH-FK-12 | 0.8 | 0.8 | 2.5 |
| JH-FK-13 | >50 | >50 | >10 |
| JH-FK-14 | 1.6 | 0.8 | 0.6 |
| JH-FK-15 | 3.1 | 0.4 | 0.3 |
| JH-FK-16 | 0.4 | 0.4 | 1.3 |
| JH-FK-17 | 0.4 | 0.4 | 1.3 |
| JH-FK-20 | 0.8 | 0.8 | 0.6 |
| JH-FK-21 | 1.6 | 0.8 | 2.5 |
| JH-FK-22 | 1.6 | 1.6 | 2.5 |
| JH-FK-23 | 1.6 | 1.6 | 5 |

<sup>a</sup> MIC<sub>90</sub> were tested at 37°C for *C. neoformans*. MIC<sub>80</sub> were tested at 30°C with 2 µg/mL supplementary fluconazole for *C. albicans*. MEC was collected for *A. fumigatus* strains.

### Modifications at C-22 position reduced inhibition of mammalian calcineurin signaling

Mammalian calcineurin inhibition was then assessed and found to be reduced for these FK520 analogs. The inhibition of mammalian calcineurin was assessed by measuring the reduction of IL-2 expression from primary mouse CD4^+^ T cells cultured in the presence of FK506, FK520, or an analog. Calcineurin activity is required for T cell activation and cytokine production, and IL-2 production serves as an effective measure of mammalian calcineurin activity. After culturing mouse CD4^+^ T cells for 72 hours, IL-2 expression was analyzed by flow cytometry to evaluate immunosuppressive activity of the analogs. The results showed an overall reduction in *in vitro* immunosuppressive activity of the analogs compared to FK506 and FK520 as denoted by the increased concentration required to inhibit cytokine production (Fig. 2A). Previous studies measuring FK506 immunosuppressive activity have demonstrated FK506 to be cytotoxic and reduce proliferation of T cells (42). In this study, decreased levels of cell proliferation were observed at high doses of compounds where IL-2 expression was decreased. However, compound concentrations that permit high IL-2 expression are less likely to result in cell death. Therefore, IC_50_ values were determined from these data to quantify the reduction in immunosuppression due to the structural modification of FK520 (Table 2). The IC_50_ values of the analogs demonstrate a minimum 65-fold reduction in mammalian calcineurin inhibition. Notably, IC_50_ values increased from 0.09 nM for FK506 up to 70.5 nM for analog JH-FK-23 and to 42.6 nM for JH-FK-08. Analog JH-FK-13 showed an IC_50_ greater than 200 nM and an MIC value greater than 50 μg/mL, indicating targeting of both mammalian and fungal calcineurin has been compromised. Our results demonstrate that structurally modified analogs have reduced capacity to inhibit mammalian calcineurin, and therefore their immunosuppressive activity has been reduced.

**Table 2.** MTherapeutic indices for FK506, FK520, and analogs.

| Compound | R | H99 MIC<sub>90</sub><br>(μg/mL) | IL-2 IC<sub>50</sub><br>(nM) | Therapeutic<br>Index |
| --- | --- | --- | --- | --- |
| FK506 | - | 0.05 | 0.09 | 1.0 |
| FK520 | - | 0.05 | 0.3 | 3.3 |
| JH-FK-08 | CO<sub>2</sub>Me | 0.8 | 42.6 | 29.6 |
| JH-FK-09 | C(=O)-4-pyridine | 0.8 | 11.2 | 7.8 |
| JH-FK-12 | CONH<sub>2</sub> | 0.8 | 35.6 | 24.7 |
| JH-FK-13 | SO<sub>2</sub>-4-Me-Ph | >50 | >200 | undefined |
| JH-FK-14 | C(=O)-2-CONHNH<sub>2</sub>-Ph | 1.6 | 14.0 | 4.9 |
| JH-FK-15 | C(=O)-2-naphthalene | 3.1 | 17.4 | 3.1 |
| JH-FK-16 | Et | 0.4 | 5.9 | 8.2 |
| JH-FK-17 | C(=O)-4-F-Ph | 0.4 | 9.4 | 13.1 |
| JH-FK-20 | C(=O)-4-NH<sub>2</sub>-Ph | 0.8 | 17.8 | 12.3 |
| JH-FK-21 | CONHNH<sub>2</sub> | 1.6 | 54.5 | 18.9 |
| JH-FK-22 | COCH<sub>2</sub>OH | 1.6 | 45.0 | 15.6 |
| JH-FK-23 | SO<sub>2</sub>Me | 1.6 | 70.5 | 24.5 |

**Fig. 2.**
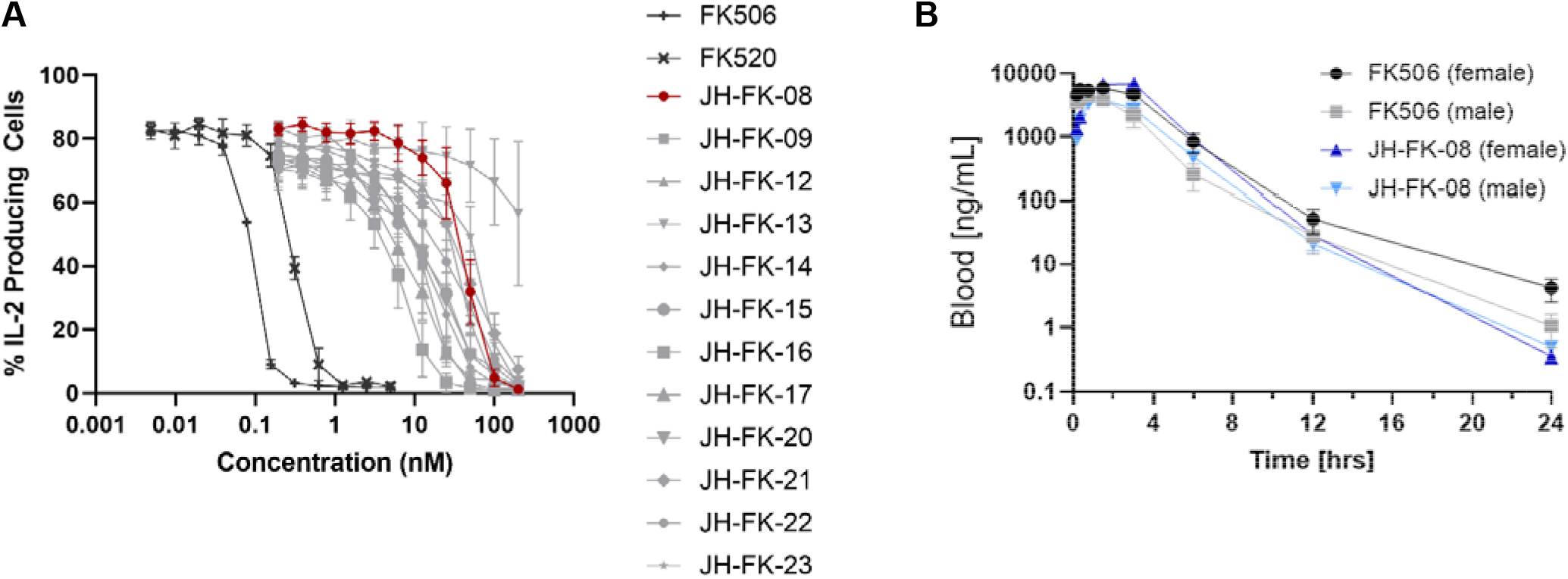
*In vitro* immunosuppressive activity of FK506, FK520, and analogs and pharmacokinetic profile of JH-FK-08 compared to FK506. (*A*) The percentage of IL-2-producing CD4^+^ T cells is plotted vs. drug concentration as a measure of immunosuppressive activity of calcineurin inhibitors. Newly synthesized analogs demonstrate reduced immunosuppression compared to controls FK506 and FK520. (n=3, error bars represent standard error of the mean (SEM), IC_50_ values determined by nonlinear regression with variable slope analysis). (*B*) Whole blood drug concentrations are plotted over time following single dose of 40 mg/kg of FK506 or JH-FK-08 (n=3, no statistically significant difference between groups, repeated measures ANOVA with Greisser Greenhouse correction test).

### JH-FK-08 has been identified as a lead compound

Based on the *in vitro* fungal and mammalian calcineurin inhibition data, the panel of FK520 analogs was assessed and ranked. Therapeutic index scores were used to quantify the effects of structural modification on the compounds (Table 2). Therapeutic index scores (TI) were calculated for each analog as the ratio of relative change in immunosuppressive and antifungal activities compared to FK506 as previously utilized (46). This ratio of therapeutic benefit and host toxicity, or fungal and mammalian calcineurin inhibition, allowed us to quantify and rank these analogs. Using the formula TI = (IC_50[compound]_/IC_50[FK506]_) / (MIC_[compound]_/MIC_[FK506]_) and the *in vitro* immunosuppressive and antifungal activities against *C. neoformans*, the TI for JH-FK-08 is 29.6. This is the greatest TI found within the library and allowed us to identify JH-FK-08 as a lead candidate for *in vivo* study. Tolerability of JH-FK-08 was assessed by monitoring animal response to drug dosing across 14 days including changes in weight, fur condition, and behavior. A maximum dose of 40 mg/kg was tested in single day dosing, and due to solubility limitations, tolerability of JH-FK-08 at 60 mg/kg and 80 mg/kg total daily dose was assessed by dosing two equal doses per day (Fig. S2). Animals tolerated all dosages, and weight ranges remained within ∼10% of vehicle treated animals. It should be noted that analogs JH-FK-12 and JH-FK-23 are also promising candidates with respective TIs of 24.7 and 24.5.

For the lead compound JH-FK-08, the mechanism of action resulting in antifungal activity was tested for conservation of calcineurin inhibition. To probe the JH-FK-08 mechanism of action, MICs of JH-FK-08 were determined with fungal mutants that lack the drug binding protein FKBP12 or have mutations in the binding site on calcineurin that prevent FKBP12-FK506 binding and inhibition. These results demonstrate: 1) FKBP12 is required for the antifungal activity of JH-FK-08, similar to the action of FK506 and FK520, 2) mutations of calcineurin B that render calcineurin resistant to binding and inhibition by FKBP12-FK506 similarly confer resistance to JH-FK-08, and 3) FK506, FK520, and JH-FK-08 exhibit potent antifungal activity against *C. neoformans* at 37°C but not at 30°C, in accord with the known requirement of calcineurin for growth at elevated temperature (Table S2). Taken together, these findings support the hypothesis that JH-FK-08 requires FKBP12 for antifungal activity, and the target of the FKBP12-JH-FK-08 complex is calcineurin. The length of time that drugs are active *in vivo* is critical information relevant to drug development, and the time frame of drug activity can have a drastic impact on treatment regimen and efficacy. Therefore, the pharmacokinetic profiles of FK506 and JH-FK-08 were investigated. Three male and three female CD1 mice were administered a single dose of JH-FK-08 or FK506 at 40 mg/kg. Whole blood samples were taken over a 24-hour period and the drug concentration remaining in the blood was determined. Pharmacokinetic parameters were calculated by a non-compartmental approach (52). No significant differences across groups were observed in whole blood samples of drug concentration over time, indicating a strong conservation of the pharmacokinetic profile of JH-FK-08 compared to that of FK506 (Fig. 2B). When assessing specific pharmacokinetic measures sex differences began to emerge. When looking at the area under the curve (AUC), females showed roughly twice the value of males, and male clearance of both FK506 and JH-FK-08 was higher compared to female treated animals (Table S3). Despite this, the half-life across all groups ranged from 1.14-1.48 hours. Overall, JH-FK-08 exhibits a pharmacokinetic profile very similar to that of FK506 including sex-dependent trends seen in clearance and AUC.

### *In vivo* immunosuppressive activity of JH-FK-08 is reduced

Whole animal studies were conducted to assess the *in vivo* implications of the reduced mammalian calcineurin inhibition demonstrated during *in vitro* immune cell testing. The *in vivo* immunosuppressive effect of JH-FK-08 was assessed by studying T follicular helper (Tfh) cell production in mice. Tfh cells play a role in protection against and response to fungal infections (53, 54). For this study, immunocompetent C57BL/6 mice received 8 days of twice daily dosing of vehicle, FK506 (2.5 mg/kg), or JH-FK-08 (20 or 40 mg/kg). Twenty-four hours after the first dose, mice were immunized with NP-OVA to induce an immune response. Cells were analyzed 7 days post immunization, and cell types were analyzed by flow cytometry (Fig. 3B).

**Fig. 3.**
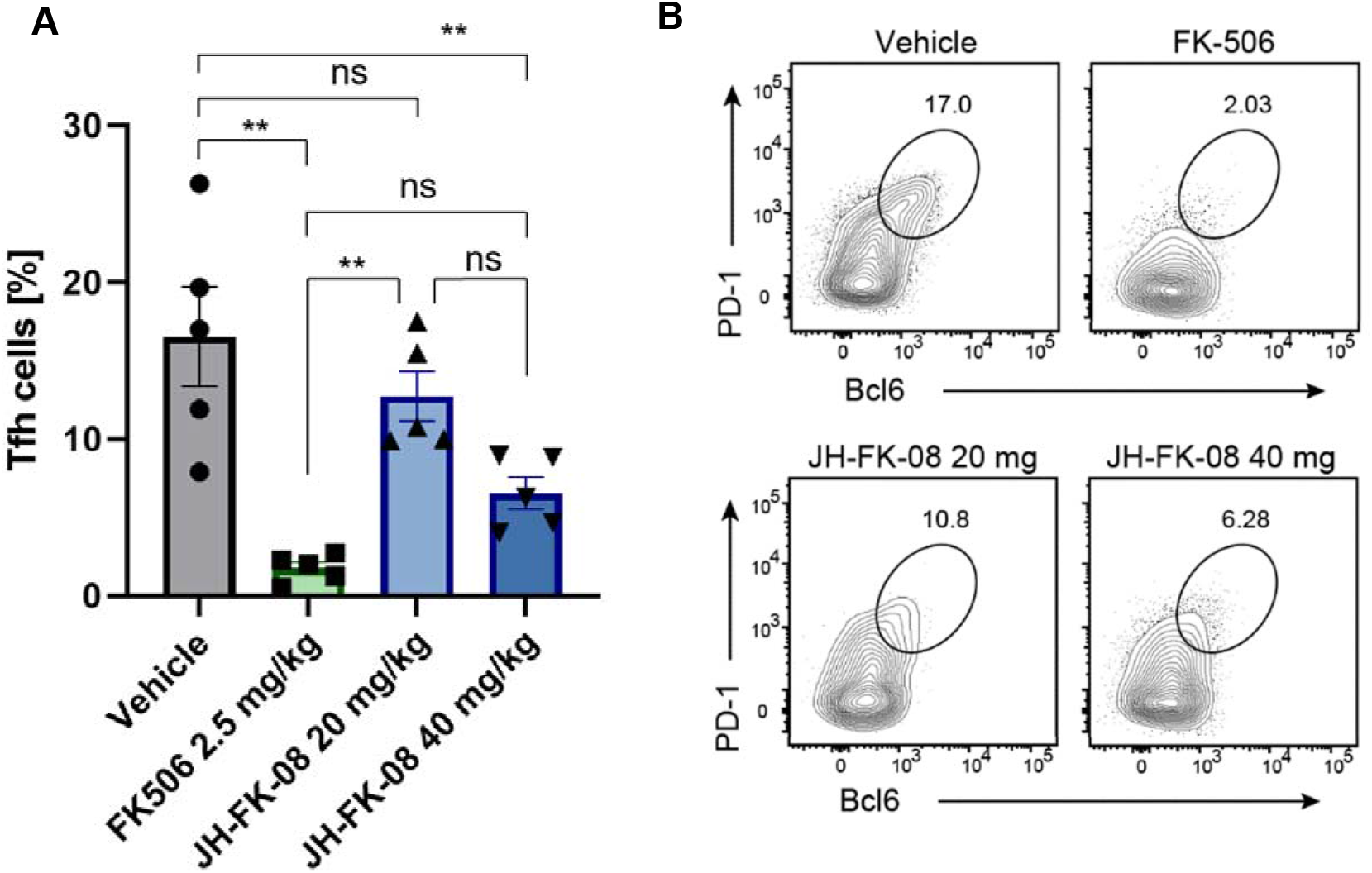
*In vivo* immunosuppression activity of JH-FK-08 is reduced compared to FK506. (*A*) Shown here are the production of T follicular helper (Tfh) cells during immune response in C57BL/6 female mice. (*B*) Shown here are representative flow cytometry plots showing the frequencies of Tfh cells among CD44^hi^ CD4^+^ T cells. Mice received twice daily treatment of vehicle, FK506 (2.5 mg/kg), JH-FK-08 (20 mg/kg), or JH-FK-08 (40 mg/kg) (n=5; **, p<0.001, ordinary One-way ANOVA with Tukey’s test).

Treatment with JH-FK-08 at 20 mg/kg had no significant effect on the percent of Tfh cells produced compared to vehicle (Fig. 3A). However, the production of Tfh cells was reduced at a dosage of 40 mg/kg of JH-FK-08. The immune response was also significantly reduced in mice treated with 2.5 mg/kg FK506. In germinal center B cells, JH-FK-08 showed similar levels of immunosuppressive activity compared to FK506 (Fig. S3). Overall, these findings indicate that Tfh cells are impacted by JH-FK-08 in a dose-dependent manner, and FK506 significantly inhibits Tfh cell production at 1/16^th^ of the dose of JH-FK-08. These results in whole animals are well correlated with the finding that the analog JH-FK-08 has significantly reduced mammalian calcineurin inhibition *in vitro* compared to FK506.

### JH-FK-08 reduces fungal burden during murine cryptococcal infection

*In vivo* antifungal activity and therapeutic benefit was assayed by investigating the impact of the FK520 analog on organ fungal burden during cryptococcosis. Specifically, *in vivo* efficacy of JH-FK-08, on fungal burden was assessed using A/J female mice. Immunodeficient A/J mice were selected in part due to their defect in complement component 5 and are susceptible to cryptococcal infection. Treatment groups included vehicle of 10% Kolliphor : ethanol (1:1) in phosphate buffered saline (PBS), 6 mg/kg fluconazole, 40 mg/kg JH-FK-08, or the fluconazole and JH-FK-08 combination, all given via intraperitoneal (i.p.) injection twice per day beginning at four hours post infection. Single day dosing with vehicle, 12 mg/kg fluconazole, 40 mg/kg JH-FK-08, and JH-FK-08 plus fluconazole combination resulted in limited efficacy reducing lung fungal burden and dosing twice per day was implemented to augment treatment efficacy (Fig. S4). Mice were intranasally infected with *C. neoformans* strain H99, and fungal burden in the lung, spleen, and brain was assessed after 14 days of treatment. Treatment with fluconazole, JH-FK-08, or a combination resulted in no detectable fungal burden in the brain or spleen. The average lung fungal burden in untreated animals was approximately 3.7 million fungal cells per gram of lung tissue. This lung fungal burden was significantly reduced by ∼6-fold, ∼200-fold, and by >400-fold in treatments with fluconazole, JH-FK-08, or the combination, respectively (Fig. 4). Additionally, recruitment of immune cells to the site of infection in lung tissue occurred based on histopathology analysis. The expected granulomatous inflammatory response admixed with mature lymphocytes and neutrophils was seen during JH-FK-08 and combination treatment (Fig. S5). This indicates that reduction of JH-FK-08 immunosuppression was sufficient such that immune responses were still mounted during infection, and JH-FK-08 treatment had a direct impact reducing organ fungal burden during cryptococcal infection.

**Fig. 4.**
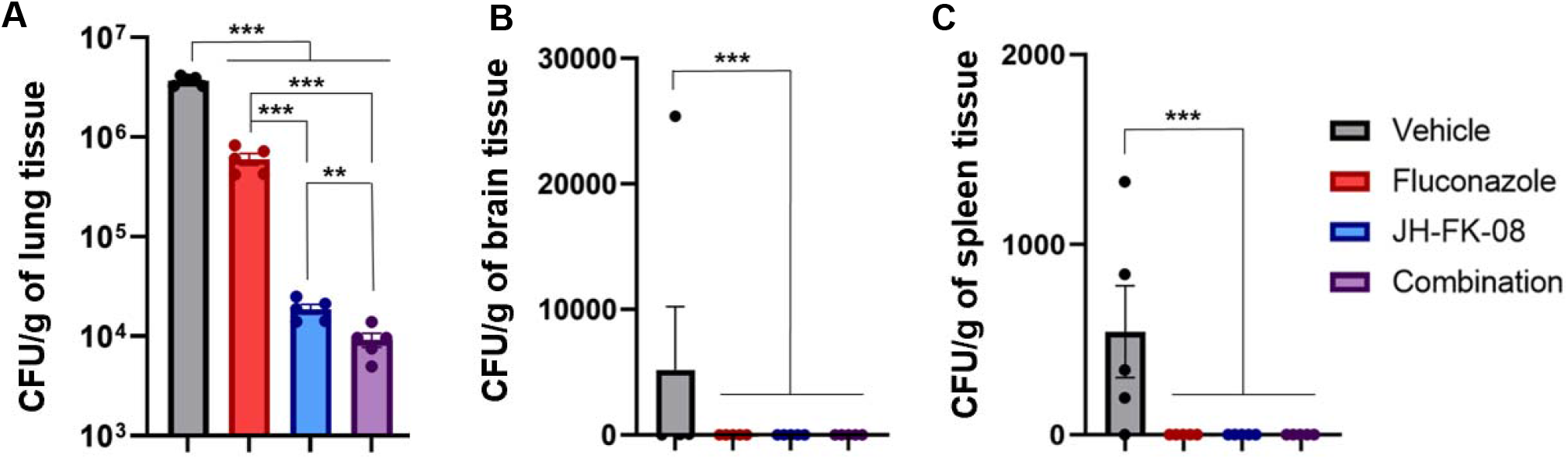
JH-FK-08 treatment significantly reduced organ fungal burden in infected animals. Female A/J mice (n=5) were infected with *C. neoformans* strain H99 through intranasal inoculation and treated twice daily with vehicle, 6 mg/kg fluconazole, 40 mg/kg JH-FK-08, or combination through i.p. dosing. Fungal burden levels in lung (*A*), brain (*B*), and spleen (*C*) were assessed after 14 days of treatment. (***, p<0.0001; **, p=0.0018, ordinary 1-way ANOVA with Dunnet’s test, error bars represent SEM)

### Treatment with JH-FK-08 extends survival during cryptococcal infection

Finally, whether the antifungal efficacy translates into extended survival during infection was assessed. To test treatment impact on survival, murine survival was monitored in the cryptococcosis infection model. Following the same protocol of infection and dosing as in the fungal burden experiment, animal survival was monitored following infection and two weeks of intraperitoneal dosing. Fluconazole treatment extended median survival from 16 to 27 days. JH-FK-08 treatment alone more than doubled median survival from 16 days in vehicle treated mice to 33 days in treated animals. This beneficial impact on survival was further enhanced by concomitant fluconazole treatment, and this resulted in an increased median survival from 16 days to 36 days (Fig. 5). These results further emphasize the therapeutic benefit JH-FK-08 offers by demonstrating an extended survival of animals treated with JH-FK-08 alone or in combination with fluconazole.

**Fig. 5.**
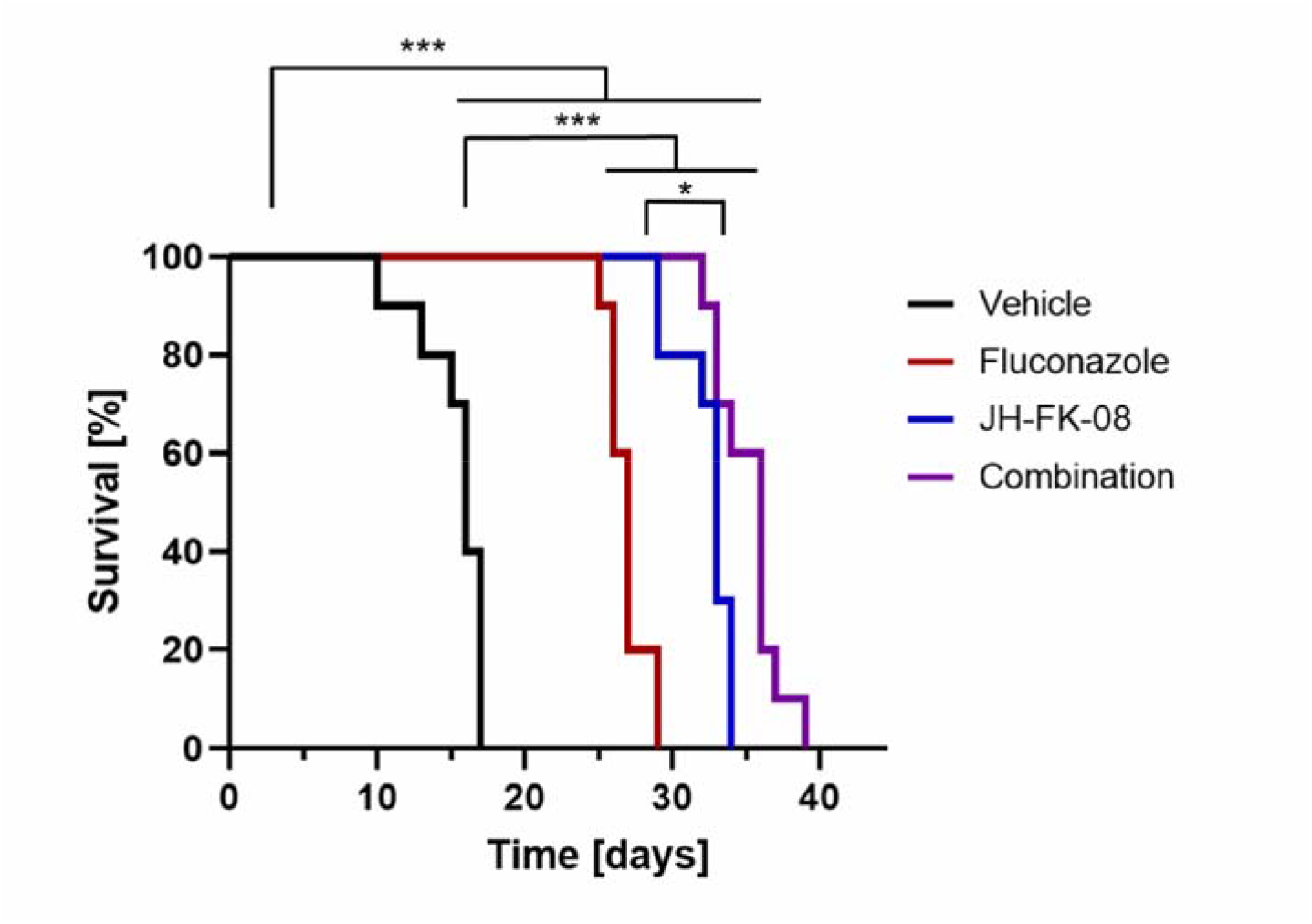
Murine survival is significantly extended by treatment with JH-FK-08 alone or in combination with fluconazole. Female A/J mice (n=10) were intranasally infected with *C. neoformans* strain H99 and received twice daily dosing of 6 mg/kg fluconazole, 40 mg/kg JH-FK-08, or a combination of the two drugs for the first two weeks post infection. Survival was monitored following the treatments (*, p=0.0065 found between JH-FK-08 and combination treatment groups; ***, p<0.0001 found between all other groups, Mantel Cox test).

## Discussion

In this study, an expanded library of structurally validated FK520 analogs was generated by one-step chemical synthesis and screened for their antifungal and immunosuppressive activity to develop non-immunosuppressive fungal-specific calcineurin inhibitors. The research and development of novel antifungal therapeutics is relevant due to rapidly emerging resistance to the current antifungal arsenal (6, 7). Additionally, cryptococcal infections are associated with high mortality, and the treatment options for the population most devastated by this human fungal pathogen are typically limited to orally available but fungistatic fluconazole (1, 3). FK506 is an orally active drug, making the development of fungal-specific analogs a potential avenue to better serve these patients.

By first assessing *in vitro* antifungal and immunosuppressive activity of the various C-22 modified FK520 analogs, we were able to confirm fungal specificity. During antifungal *in vitro* screening, fungal calcineurin inhibition was measured through microdilution broth growth assays. These results demonstrated many compounds were able to maintain potent antifungal activity indicating these structural modifications did not prevent fungal calcineurin targeting. Reduced mammalian calcineurin inhibition by the FK520 analogs was confirmed in cytokine production studies. During this *in vitro* immunosuppressive assay, production of the cytokine IL-2, which is known to be inhibited by FK520, was shown to be suppressed to a lesser degree by the FK520 analogs. This finding confirms reduced targeting of the mammalian calcineurin by the FK520 analogs. Using this information, therapeutic index scores were calculated to quantify the degree to which fungal selectivity was achieved.

JH-FK-08 was identified as the lead compound following *in vitro* screening. The introduced structural modifications did not significantly alter the pharmacokinetic (PK) properties (e.g., C_max_, half-life, whole-blood drug concentration over time) compared to FK506 in both male and female mice. A pattern of higher plasma exposure (higher AUC, decreased clearance) in females was also observed in JH-FK-08- and FK506-treated animals relative to male treated animals (Table S3). To further study the antifungal effect of calcineurin inhibition during infection, JH-FK-08 was then investigated for *in vivo* antifungal efficacy. Animals were intranasally infected with *C. neoformans* to replicate the conventional infection route in humans. In this study, JH-FK-08 was found to significantly reduce lung fungal burden by approximately 200-fold during two weeks of treatment and more than doubled murine survival time during infection compared to untreated infected animals. These benefits were further enhanced when animals were treated with JH-FK-08 in combination with fluconazole.

During this study, dosing animals with either JH-FK-08 alone or in combination with fluconazole significantly reduced fungal burden and extended survival during cryptococcal infection. However, during this treatment time frame, infection was not fully cleared, and therefore infection progression led to mortality even in treated animals. One potential future direction is to extend the treatment time to assess if clearance can be achieved.

Overall, the preclinical efforts presented here advance JH-FK-08 as a promising candidate for potential clinical study. The activity of JH-FK-08 far exceeds the previous two libraries’ top candidates APX879 and JH-FK-05 based on therapeutic index score comparison and *in vivo* efficacy (44, 45). These results support further study of calcineurin inhibition as a potential effective antifungal approach. Exciting research avenues for future study include: 1) investigating oral activity of JH-FK-08, 2) researching how JH-FK-08 can interact with and cross the blood brain barrier, and 3) investigating possible combination therapies. Current treatments for fungal infections make use of a combinations of therapies, and using multiple drugs during one therapeutic application can increase treatment efficacy as well as reduce emerging resistance and mitigate potential side effects (5, 49). One potential combination partner is APX001A and its analogs as this compound is under clinical investigation as a novel antifungal class (47, 50, 51). Furthermore, this approach of designing fungal specific analogs of existing therapies can be utilized with other natural FKBP12 binding products such as rapamycin, which also demonstrates potent antifungal activity against high priority fungal pathogens (52, 53).

### Materials and methods Synthesis of FK520 analogs

General procedure A: To a solution of FK520 (APIChem Technology) in EtOH (0.15 M) was added a hydrazide (6 eq) in a sealed vial. The reaction mixture was stirred at 90 °C for 48 h. After cooling to 25 °C, the reaction mixture was concentrated *in vacuo*. The crude mixture was reconstituted in EtOAc and washed with H_2_O and brine. The combined organic layers were dried over anhydrous Na_2_SO_4_ and concentrated *in vacuo*. The residue was purified by column chromatography (silica gel, CH_2_Cl_2_/MeOH) to afford a substituted hydrazone analog.

General procedure B: To a solution of FK520 in MeOH (0.10 M) was added a hydrazide (6 eq) or a substituted hydrazine (6 eq) and TFA (1 μL per 100 mg of FK520). The reaction was stirred at 25 °C for 48 h. The reaction mixture was concentrated *in vacuo* and reconstituted in EtOAC. The resulting reaction mixture was washed with 10% aqueous NaHCO_3_ solution, H_2_O, and brine. The combined organic layers were dried over anhydrous Na_2_SO_4_ and concentrated *in vacuo*. The residue was purified by column chromatography (silica gel, CH_2_Cl_2_/MeOH) to afford a substituted hydrazone analog.

### Antifungal susceptibility testing

*In vitro* antifungal activity was assessed for all fungal strains in RPMI 1640 (Sigma-Aldrich R1383). MICs were measured using Clinical and Laboratory Standards Institute (CLSI) M38-A2 and M27-A3 standard *in vitro* antifungal susceptibility protocols. *C. neoformans* and *A. fumigatus* testing was performed at 37°C. MECs for *A. fumigatus* were recorded after 48 h growth. *C. albicans* testing was performed at 30°C with a uniform concentration of 2 μg/mL of fluconazole to sensitize the cells to calcineurin inhibition.

### Fungal strains

*C. neoformans* clinical isolate H99 and congenic derived strain KN99α strain were used (55). KN99α derived *frr1*Δ deletion mutant with nourseothricin resistance marker as part of the Madhani deletion collection (NIH funding R01AI100272) was used. *C. deneoformans* strains used include JEC21 and derived C21F2 and C21F3 strains (20, 56, 57). *C. albicans* strain SC5314 is the wild type strain from which mutants YAG171 (*rbp1/rbp1*) and YAG237 (*CNB1-1*/*CNB1*) were derived (58). *A. fumigatus* strain *akuB*^Ku80^ and a derived FKBP12 mutant *fkbp12*Δ were used during antifungal screening (59).

### *In vitro* immunosuppressive activity testing

Spleen and lymph nodes from C57BL/6 mice were collected and homogenized through a 40 μm filter. Pan-CD4^+^ T cells were enriched using MagniSort Mouse CD4 T cell Enrichment Kit (Invitrogen). Thereafter, CD4^+^ CD25^-^ CD62L^hi^ CD44^lo^ naïve CD4^+^ T cells were FACS-sorted using a MoFlo Astrios cell sorter (Beckman Coulter). Naïve CD4^+^ T cells were cultured in Iscove’s Modified Dulbecco’s Medium (IMDM) supplemented with 10% FBS, penicillin (10LJU/ml), streptomycin (10LJμg/ml), glutamine (2 mM), gentamicin (50 μg/ml), and β-mercaptoethanol (55 μM). Cells were cultured for 72 h on anti-hamster IgG-coated plates with hamster anti-CD3ε antibodies, hamster anti-CD28 antibodies, neutralizing anti-IL-4 antibody, recombinant IL-12 (10 ng/mL), and recombinant IL-2 (50 U/mL). During this culture, cells were incubated with serially diluted drug suspended in DMSO.

### Flow cytometry

Cell surface staining of single cell suspension was performed in staining buffer (PBS supplemented with 2 mM EDTA and 0.5% BSA). Anti-FcγR (93; Biolegend) was treated to block Fc receptor before surface staining of primary cells isolated from lymph nodes of NP-OVA-immunized mice. Cells were stained with fixable viability dye (eBioscience) before surface staining to exclude dead cells from analysis. For the detection of intracellular cytokines, cells were additionally incubated with phorbol 12-myristate 13-acetate (PMA), ionomycin in the presence of GolgiStop (BD Biosciences) during the final 4 h of incubation. For intracellular staining, cells were fixed and permeabilized with Fixation/Permeabilization buffer (eBioscience). Flow cytometric analysis was performed utilizing BD FACSCantoII or BD LSRFortessaX20 (BD Biosciences) and FlowJo software (BD Biosciences).

### Compound preparation for *in vivo* studies

For *in vivo* studies, fluconazole was purchased from Duke Pharmacy Storeroom. Vehicle was a sterile solution of 10% Kolliphor EL (Sigma-Aldrich C5315), 10% ethanol, and 80% phosphate-buffered saline (PBS, Sigma-Aldrich D8537). For administration to animals, all other drugs were obtained in powder form and diluted in a sterile mixture of 50% Kolliphor EL 50% ethanol before dilution to 80% PBS.

### *In vivo* immunosuppressive assessment

Groups of 5 female C57BL/6 mice were treated twice daily for 8 days with vehicle, FK506 (2.5 mg/kg), JH-FK-08 (20mg/kg), or JH-FK-08 (40 mg/kg) via intraperitoneal (i.p.) injection. 24 h after the first dose, animals were immunized with NP-OVA (100 μg) in alum via subcutaneous injection. 7 days post-immunization, mice were sacrificed, and draining lymph nodes were harvested for the analysis of Tfh cells and GC B cells by flow cytometry.

### *C. neoformans* murine infection

*C. neoformans* inoculum was prepared by growing a 5 ml overnight YPD culture spinning in a tissue culture roller drum at 30°C. Cells were pelleted by centrifugation and washed in sterile PBS 3 times, concentration was determined by hemocytometer, and cells were diluted to 2x10^6^ cells/mL. Mice (10 to 20 g) were infected with 10^5^ cells of *C. neoformans* H99. For inhalation infection, animals were anesthetized with isoflurane before 50 μl of inoculum was pipetted onto the upright nasal passages of each animal. Following inhalation of the inoculum, animals were monitored for post anesthesia recovery. Treatment of animals began 4 hours post infection and was continued for 14 days. Fungal burden was assessed 24 hours after the final dose.

### Histology preparation

Lungs were inflated with and immersed in 10% neutral buffered formalin, processed routinely to paraffin embedment and sectioned at 5 mm. Sections were stained either with Hematoxylin & Eosin (H&E) or Periodic-acid Schiff (PAS). All tissue sections were evaluated and photographed with an Olympus BX51 microscope with a DP70 camera using Olympus DP Manager software v.3.3.112. Samples processed by the Department of Pathology.

### Pharmacokinetics (PK) experiment

Male (n=3; avg. 28 g) and female (n=3, avg. 25 g) CD-1 mice were injected intraperitoneally with 40 mg/kg of either JH-FK-08 or FK506. The drug formulation was 10% Kolliphor/10% ethanol/80% PBS given as 10 μl/g body weight (i.e., 200 μl per 20 g mouse). Blood (∼30 μl) was collected serially (“tail snip”) at 10, 20, 45 min, 1.5, 3, 6, 12, and 24 h into vials containing 1 ml of 75 mg/ml K_2_EDTA in water, and immediately frozen until the day of LC/MS/MS analysis. Blood concentration/time data was utilized to calculate relevant PK parameters by non-compartmental approach and extravascular administration model within WinNonlin (2.1) software.

### LC/MS/MS analysis

In 0.5 ml polypropylene vial, 10 μl blood and 20 μl of acetonitrile fortified with 100 ng/ml FK520 (internal standard), were vigorously agitated in FastPrep FP120 apparatus (Thermo-Savant) at speed 4 for 45 s. After centrifugation for 5 min at 16,000 rcf, 20 μl of supernatant was transferred to polypropylene autosampler vial. Liquid chromatography - tandem-mass spectroscopy (LC/MS/MS; Agilent 1200 series HPLC and Sciex/Applied Biosystems API 5500 QTrap) was utilized to quantify JH-FK-08 and FK506 in mouse blood. Analytical column: Agilent Eclipse Plus (C18, 1.8 μm, 50 × 4.6 mm), at 40°C. Mobile phase: (A) 0.1% formic acid, 10 mM ammonium acetate, 2% acetonitrile in water, (B) methanol. Elution gradient: 0-0.1 min 30-98% B, 0.1-1.5 min 98% B, 1.5-1.6 min 98-30% B, 1.6 -2.1 min 30% B. Run time: 5 min. Injection volume: 10 μl. Mass spectrometer parameters (voltages, gas flow, and temperature) were optimized by infusion of 100 ng/mL of analytes in mobile phase at 10 μl/min using Analyst 1.6.2 software tuning module. The MS/MS (m/z) transitions used for quantification: 864.4/846.5 (JH-FK-08), 821.4/768.5 (FK506), and 809.4/756.5 (FK520, internal standard). A set of calibrator samples in drug-free blood was prepared by adding appropriate amounts of pure analyte (JH-FK-08 or FK506) in 2.43 - 1000 μl/ml range. The calibration samples were analyzed alongside the experimental samples. Accuracy acceptance criteria was 85% for each but the lowest level (2.43 ng/mL, 80%, LLOQ). Analyst 1.6.2 software was used for data acquisition, integration of the chromatograms, calibration curve calculation, and quantification of the study samples.

### *Cryptococcus* fungal burden and survival analysis

For survival curve analysis, animals were monitored until reaching defined humane end points, including social isolation, hunched posture, weight loss, lack of grooming, and labored breathing, and were then euthanized via CO_2_ inhalation. For fungal burden analysis, mice were euthanized following 2-weeks of treatment with designated antifungal, vehicle, or combination. Brain, lung, and spleen were harvested from animals, weighed, and homogenized in sterile PBS using steel beads and a bead beater. Each organ homogenate was serial diluted 10-fold to 10^-4^ in PBS. 100 μl of each dilution was plated to YPD plates supplemented with YPD 50 μg/ml Ampicillin, 30 μg/ml chloramphenicol and incubated at 30°C for 2 days and assessed for colony forming units. CFU per organ weight was calculated and plotted using GraphPad Prism 9.

## Acknowledgements

We thank Catherine Denning-Jannace and Jeffrey Everitt for their expertise and assistance. I.S. (PK/PD Core Laboratory) acknowledges support of the NCI/NIH Comprehensive Cancer Center Core Grant P30CA014236. M.C. acknowledges support for R01 GM115474. This material is based upon work supported by the National Science Foundation Graduate Research Fellowship under Grant No. DGF 2139754. This research is supported by NIH/NIAID R01 grant AI172451-01 and R56 grant AI112595-05 and the Gilhuly Accelerator Fund.

Joseph Heitman is co-director and fellow of the CIFAR Program Fungal Kingdom: Threats & Opportunities.

## Ethics statement

Experiments utilizing mice were conducted in compliance with guidelines issued by the US Animal Welfare Act and by the Duke Institutional Animal Care and Use Committee (IACUC). Duke’s Animal Care and Use Program has been accreditation by the Association for Assessment and Accreditation of Laboratory Animal Care (AAALAC), and its animal care and housing facilities are maintained by Duke’s Division of Laboratory Animal Resources (DLAR). All procedures involving animals were approved by Duke IACUC under protocol no. A098-22-05.

## Notes

### Competing Interest Statement

The authors have declared no competing interest.

### Summary of Updates

In response, we have added additional details regarding treatment timeframe in both the main manuscript and in the methodology section. Specifically, animals began receiving treatment 4 hours post infection. Treatment continued for 14 days and fungal burden was assessed 24 hours following the final dose. Thank you for highlighting this. In response, we have edited the language on page 8 from confirming the target of the FKBP12-JH-FK-08 complex to instead supporting the hypothesis FKBP12 is required for JH-FK-08 efficacy and that the complex targets calcineurin. 'Through in vitro antifungal and immunosuppressive testing of analogs of FK520 (natural analog of FK506) analogs' has been changed to 'Through in vitro antifungal and immunosuppressive testing of FK520 (a natural analog of FK506) derivatives' Thank you for this comment. In response, we have provided the following rationale for the condensation reaction on page 6: 'To selectively modify the C-22 position on FK520, we explored the condensation reaction with hydrazine derivatives. The condensation reaction with hydrazine derivatives exploits the unique reactivity of carbonyl groups, such as ketones and aldehydes, towards hydrazine derivatives in the presence of other functional groups.' 'Modifications at the C-22 position lead to enhanced fungal specificity' has been changed to 'Modifications at C-22 position reduced inhibition of mammalian calcineurin signaling.' Further, we have included rationale behind the approach in the introduction and viability commentary in the results. We have included tolerability data in the supplemental information and expanded within the text the details regarding the tolerability study of JH-FK-08. We have included a relevant citation for the non-compartmental pharmacokinetic analysis. Great question. For Figure 3, we utilized C57BL/6 mice because they are fully immunocompetent. This characteristic allows us to elucidate the immunosuppressive impact FK506 and analogs have on a functioning immune system. However, when performing animal experiments monitoring cryptococcal susceptibility, A/J animals are used due to their immunocompromised phenotype. A/J animals are susceptible to cryptococcal infection due to their loss of function of complement component 5 and are therefore the mouse strain we selected for Cryptococcus infection models. We have included We have added additional details within the text. Although we do not have fungal burden data for animals in the survival study, we have included discussion on disease progression in animals in the survival study to address this point.

