## Supplementary Information for "Enhanced fungal specificity and *in vivo* therapeutic efficacy of a C-22 modified FK520 analog against *C. neoformans*"

**Supplemental Materials**

**Text S1.** Synthesis of FK520 analogs

**General Chemistry Procedures.** All reactions were conducted in oven-dried glassware. Unless otherwise stated, all reagents were purchased from commercial suppliers and used without further purification. FK520 was purchased from APIChem Technology and used without further purification. All solvents were American Chemical Society (ACS) grade or better and used without further purification. Analytical thin layer chromatography (TLC) was performed with glass backed silica gel (60 Å) plates with fluorescent indication (Whatman). Visualization was accomplished by UV irradiation at 254 nm and/or by staining with *p*-anisaldehyde solution. Flash column chromatography was performed by using silica gel (particle size 230−400 mesh, 60 Å). All ^1^H spectra were recorded with a Bruker 500 (500 MHz) spectrometer. All ^1^H NMR δ values are given in parts per million (ppm) and are referenced to the residual isotopomer solvent signals (acetone-*d_6_*: δ = 2.05 ppm). Coupling constants (*J*) are given in Hertz (Hz) and multiplicities are indicated using the conventional abbreviation (s = singlet, d = doublet, dd = doublet of doublets, td = triplet of doublets, qd = quartet of doublets, t = triplet, q = quartet, m = multiplet or overlap of non-equivalent resonances, br = broad). Electrospray ionization (ESI) high-resolution mass spectrometry (HRMS) was recorded with an Agilent 6624 series (LC/MS-TOF trap) spectrometer to obtain the molecular masses of compounds.

**JH-FK-08**: [Method #1] JH-FK-08 was synthesized from FK520 (50 mg, 0.065 mmol) and methyl carbazate (6 eq, 34.1 mg, 0.38 mmol) in EtOH (0.4 mL, 0.15 M) according to General Procedure A. Purification by column chromatography (silica gel, CH_2_Cl_2_/MeOH, 25/1) afforded JH-FK-08 (28 mg, 51%) as a yellow solid; [Method #2] JH-FK-08 was synthesized from FK520 (100 mg, 0.13 mmol) and methyl carbazate (6 eq, 68.2 mg, 0.76 mmol) in MeOH (1.3 mL, 0.1 M) according to General Procedure B. Purification by column chromatography (silica gel, CH_2_Cl_2_/MeOH, 25/1) afforded JH-FK-08 (92 mg, 85%) as a yellow solid: ^1^H NMR (500 MHz, acetone-*d_6_*): *δ* 9.99 (s, 1H), 8.65 (s, 1H), 5.47 (s, 1H), 5.25 (s, 1H), 5.14 (t, *J* = 7.5 Hz, 2H), 4.35 (s, 1H), 4.33 (d, *J* = 17 Hz, 1H), 4.27 (d, *J* = 4.5 Hz, 1H), 3.65 (s, 3H), 3.63 (s, 3H), 3.45 (dd, *J* = 11, 15 Hz, 2H), 3.36 (s, 5H), 3.33 (s, 3H), 3.22 (q, *J* = 7.0 Hz, 1H), 3.01–2.94 (m, 2H), 2.35 (d, *J* = 16 Hz, 2H), 2.29–2.22 (m, 2H), 2.17 (d, *J* = 14 Hz, 1H), 2.13–2.10 (m, 4H), 2.03–1.98 (m, 1H), 1.92 (s, 1H), 1.90 (s, 2H), 1.81–1.74 (m, 2H), 1.66 (s, 3H), 1.65 (s, 3H), 1.61–1.55 (m, 3H), 1.53–1.47 (m, 2H), 1.40–1.29 (m, 4H), 1.11-1.08 (m, 1H), 1.05–1.03 (m, 1H), 1.02 (d, *J* = 9.0 Hz, 3H), 0.97 (d, *J* = 7.5 Hz, 3H), 0.94–0.90 (m, 1H), 0.86 (t, *J* = 7.5 Hz, 3H), 0.85 (t, *J* = 6.5 Hz, 3H).

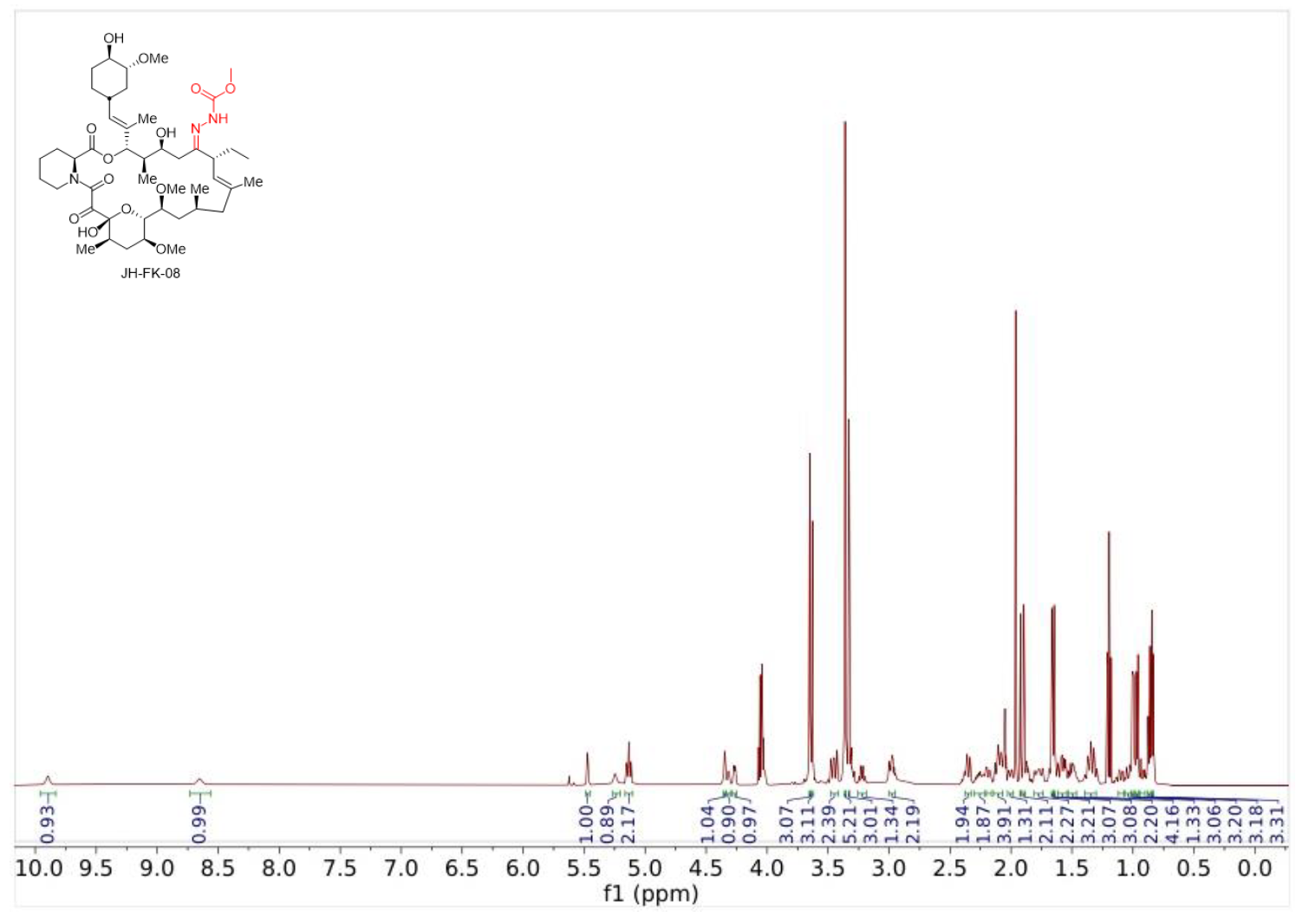

**JH-FK-09**: JH-FK-09 was synthesized from FK520 (50 mg, 0.065 mmol) and isonicotinic acid hydrazide (6 eq, 52 mg, 0.38 mmol) in EtOH (0.4 mL, 0.15 M) according to General Procedure A. Purification by column chromatography (silica gel, CH_2_Cl_2_/MeOH, 25/1) afforded JH-FK-09 (25 mg, 49%) as a white solid: ^1^H NMR (500 MHz, acetone-*d_6_*): δ 11.62 (s, 1H), 8.68 (d, *J* = 6.0 Hz, 2H), 7.76 (d, *J* = 6.0 Hz, 2H), 5.89 (s, 1H), 5.71 (s, 1H), 5.18 (d, *J* = 9.0 Hz, 1H), 5.14 (d, *J* = 9.0 Hz, 1H), 4.36 (s, 1H), 4.30–4.25 (m, 2H), 4.16 (d, *J* = 10 Hz, 1H), 3.47 (d, *J* = 10 Hz, 1H), 3.41 (t, *J* = 9.5 Hz, 1H), 3.37 (s, 2H), 3.36 (s, 2H), 3.33 (s, 2H), 3.31 (s, 5H), 3.01–2.96 (m, 2H), 2.91 (t, *J* = 13 Hz, 2H), 2.51 (d, *J* = 14 Hz, 1H), 2.42–2.34 (m, 1H), 2.23–2.09 (m, 6H), 2.04–2.00 (m, 1H), 1.96–1.85 (m, 3H), 1.81–1.72 (m, 4H), 1.69 (s, 6H), 1.64–1.54 (m, 4H), 1.37–1.28 (m, 4H), 1.10 (qd, *J =* 3.5, 12 Hz, 2H), 1.02–0.99 (m, 6H), 0.97–0.94 (m, 1H), 0.90 (t, *J* = 7.0 Hz, 3H), 0.78 (d, *J* = 6.5 Hz, 2H).

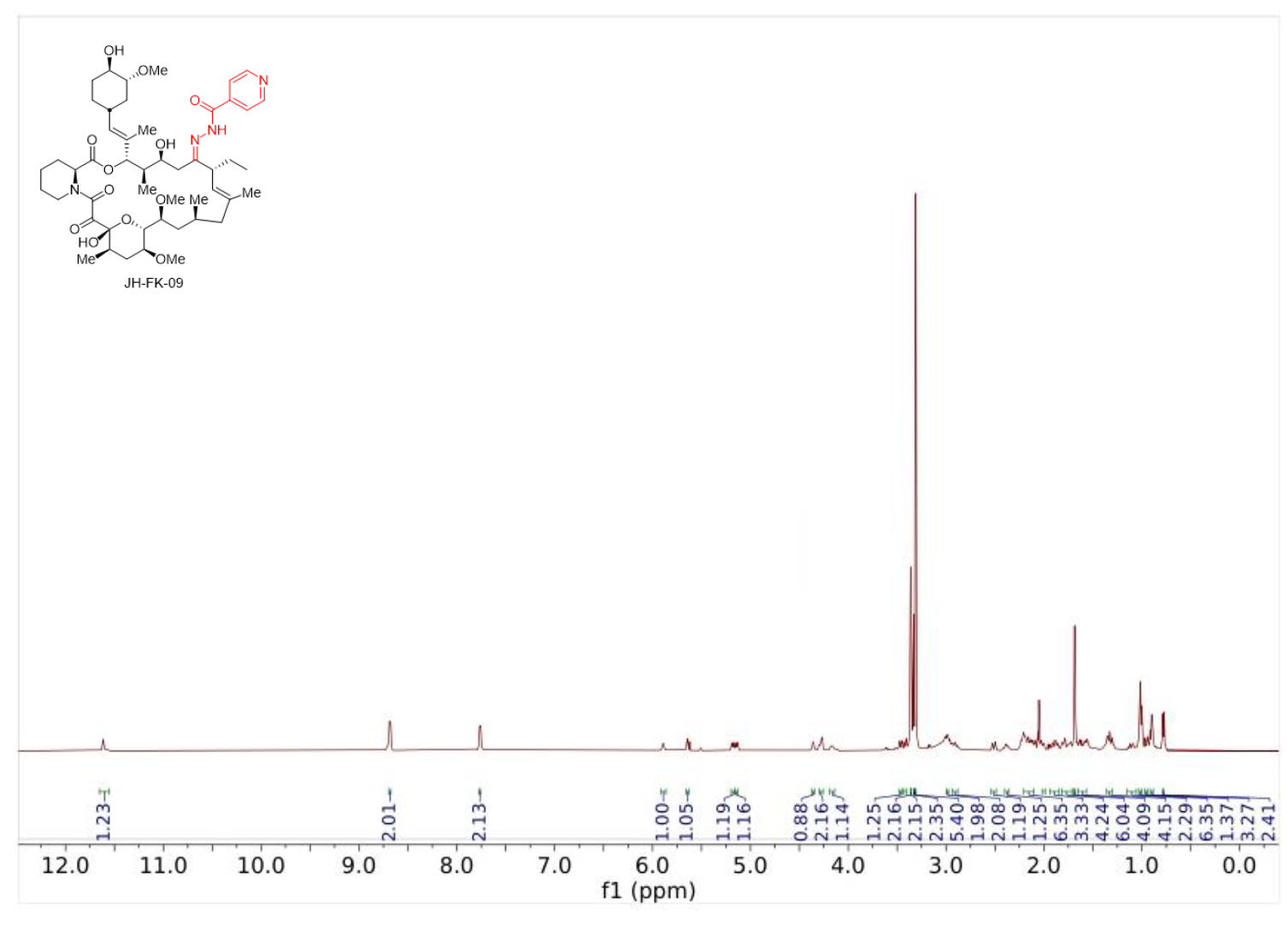

**JH-FK-12**: JH-FK-12 was synthesized from FK520 (100 mg, 0.13 mmol) and semicarbazide (6 eq, 56.8 mg, 0.76 mmol) in MeOH (1.3 mL, 0.1 M) according to General Procedure B. Purification by column chromatography (silica gel, CH_2_Cl_2_/MeOH, 25/1) afforded JH-FK-12 (41 mg, 38%) as a white solid: ^1^H NMR (500 MHz, acetone-*d_6_*): δ 9.33 (s, 1H), 6.07 (br s, 2H), 5.30 (d, *J* = 6.5 Hz, 1H), 5.22 (d, *J* = 9.5 Hz, 1H), 5.13 (d, *J* = 9.5 Hz, 1H), 4.74 (s, 2H), 4.53 (s, 1H), 4.36 (d, *J* = 14 Hz, 1H), 3.96 (s, 1H), 3.68–3.58 (m, 4H), 3.51–3.42 (m, 2H), 3.39 (s, 3H), 3.37 (s, 3H), 3.36 (s, 3H), 3.34–3.31 (m, 4H), 3.19–3.12 (m, 3H), 3.01–2.96 (m, 1H), 2.41–2.31 (m, 3H), 2.25 (s, 2H), 2.21–2.12 (m, 2H), 1.98 (s, 2H), 1.93–1.87 (m, 2H), 1.78 (s, 2H), 1.69 (s, 4H), 1.62 (s, 3H), 1.44–1.30 (m, 6H), 0.99–0.94 (m, 6H), 0.92–0.86 (m, 6H).

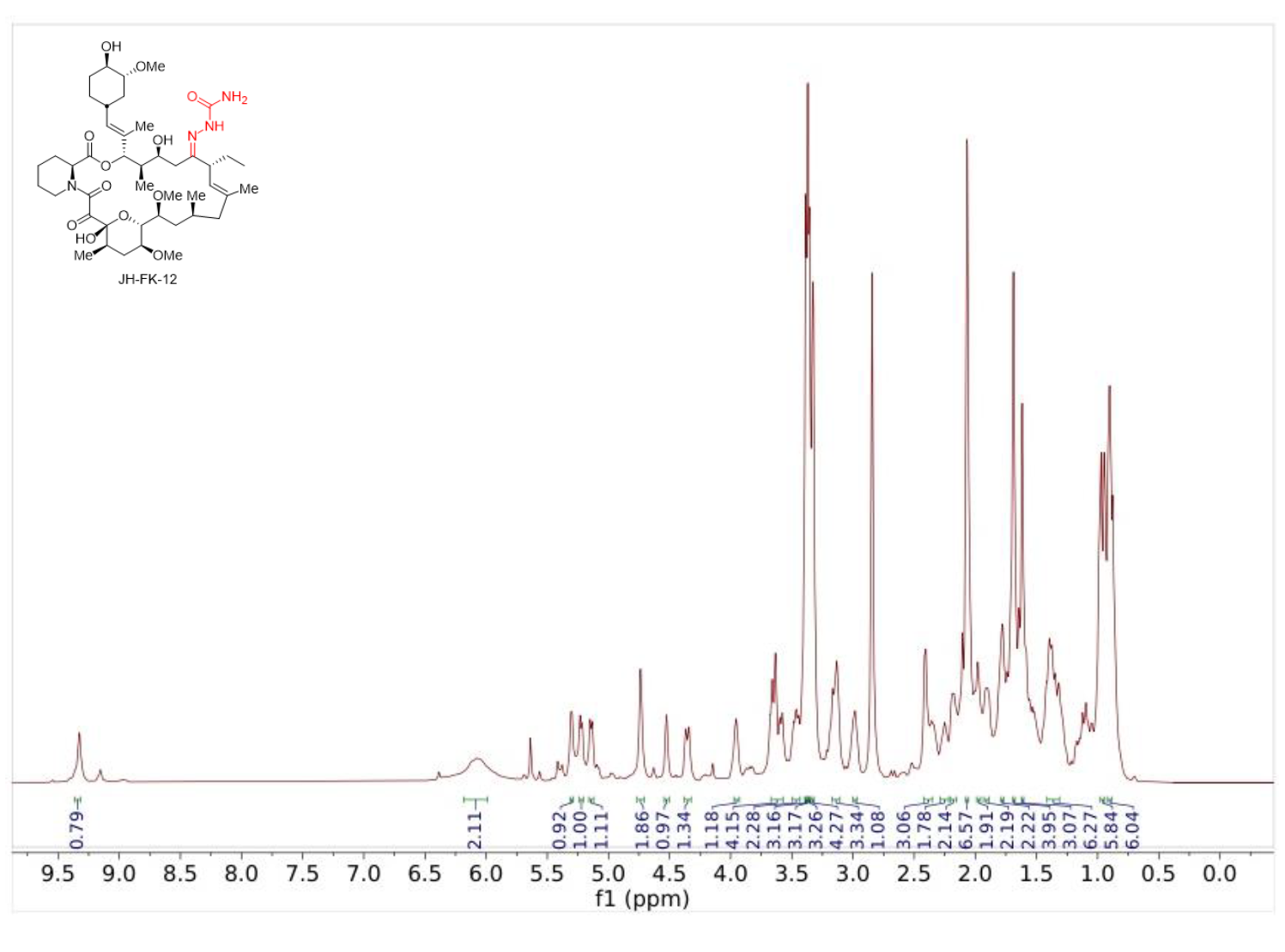

**JH-FK-13**: JH-FK-13 was synthesized from FK520 (100 mg, 0.13 mmol) and tosylhydrazine (6 eq, 141 mg, 0.76 mmol) in MeOH (1.3 mL, 0.1 M) according to General Procedure B. Purification by column chromatography (silica gel, CH_2_Cl_2_/MeOH, 25/1) afforded JH-FK-13 (40 mg, 33%) as a white solid: ^1^H NMR (500 MHz, acetone-*d_6_*): δ 9.83 (s, 1H), 7.79 (d, *J* = 8.5 Hz, 2H), 7.36 (d, *J* =11 Hz, 2H), 5.35 (d, *J* = 3.5 Hz, 1H), 5.15 (d, *J* = 9.0 Hz, 1H), 5.06 (d, *J* = 9.0 Hz, 2H), 4.60 (s, 1H), 4.40 (d, *J* = 6.0 Hz, 1H), 4.34 (d, *J* = 12 Hz, 1H), 3.91 (d, *J* = 11 Hz, 1H), 3.63 (d, *J* = 2.5 Hz, 1H), 3.52 (d, *J* = 6.5 Hz, 1H), 3.50 (d, *J* = 6.5 Hz, 1H), 3.37 (s, 2H), 3.35 (s, 2H), 3.33 (s, 2H), 3.32–3.30 (m, 1H), 3.12–3.08 (m, 2H), 3.07–2.95 (m, 2H), 2.43 (s, 1H), 2.41 (s, 2H), 2.38–2.32 (m, 1H), 2.31 (dd, *J* = 2.5, 14 Hz, 1H), 2.28–2.20 (m, 2H), 2.19–2.10 (m, 3H), 2.09 (s, 2H), 2.00–1.94 (m, 2H), 1.91–1.85 (m, 2H), 1.82–1.71 (m, 2H), 1.66 (d, *J* = 16 Hz, 1H), 1.63 (s, 3H), 1.61 (s, 1H), 1.57 (s, 3H), 1.43–1.30 (m, 7H), 1.13–1.06 (m, 2H), 0.96 (d, *J* = 6.5 Hz, 4H), 0.91 (d, *J* = 6.5 Hz, 3H), 0.89–0.85 (m, 2H), 0.83 (d, *J* = 7.0 Hz, 2H), 0.71 (t, *J* = 7.0 Hz, 1H), 0.62 (t, *J =* 7.5 Hz, 3H).

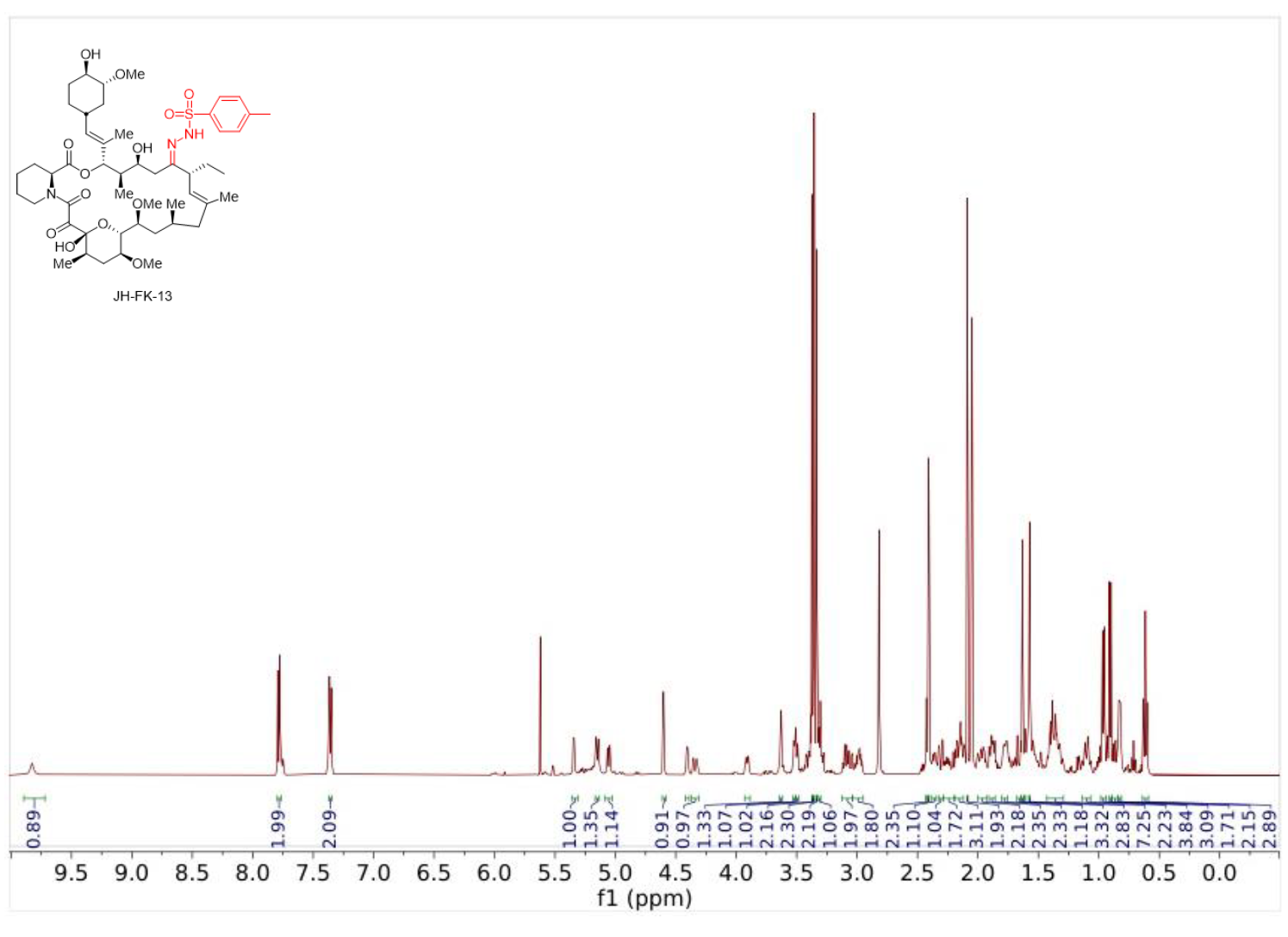

**JH-FK-14**: JH-FK-14 was synthesized from FK520 (100 mg, 0.13 mmol) and isophthalic hydrazide (6 eq, 147 mg, 0.76 mmol) in MeOH (1.3 mL, 0.1 M) according to General Procedure B. Purification by column chromatography (silica gel, CH_2_Cl_2_/MeOH, 25/1) afforded JH-FK-14 (45 mg, 37%) as a white solid: ^1^H NMR (500 MHz, acetone-*d_6_*): δ 11.76 (s, 1H), 10.04 (s, 1H), 8.30 (s, 1H), 7.98 (d, *J* = 7.5 Hz, 1H), 7.93 (d, *J* = 8.0 Hz, 1H), 7.47 (t, *J* = 8.0 Hz, 1 H), 6.19 (br s, 1H), 5.61 (s, 1H), 5.19 (d, *J* = 9.0 Hz, 1H), 5.13 (d, *J* = 9.0 Hz, 1H), 4.48 (s, 1H), 4.28–4.18 (m, 3H), 3.71 (br s, 1H), 3.45 (t, *J* = 10 Hz, 2H), 3.36 (d, *J* = 3.0 Hz, 6H), 3.33 (s, 2H), 3.31 (s, 1H), 3.00–2.95 (m, 2H), 2.94–2.88 (m, 4H), 2.51 (d, *J* = 14 Hz, 1H), 2.40–2.32 (m, 1H), 2.27–2.07 (m, 9H), 2.03–1.97 (m, 1H), 1.93 (t, *J* = 12 Hz, 1H), 1.88–1.84 (m, 1H), 1.82–1.76 (m, 2H), 1.71 (s, 3H), 1.67 (s, 3H), 1.58–1.52 (m, 2H), 1.39–1.26 (m, 5H), 1.12–1.05 (m, 2H), 1.02 (t, *J* = 10 Hz, 3H), 1.00 (t, *J* = 7.5 Hz, 4H), 0.93 (m, 4H), 0.81 (d, *J* = 6.5 Hz, 3H).

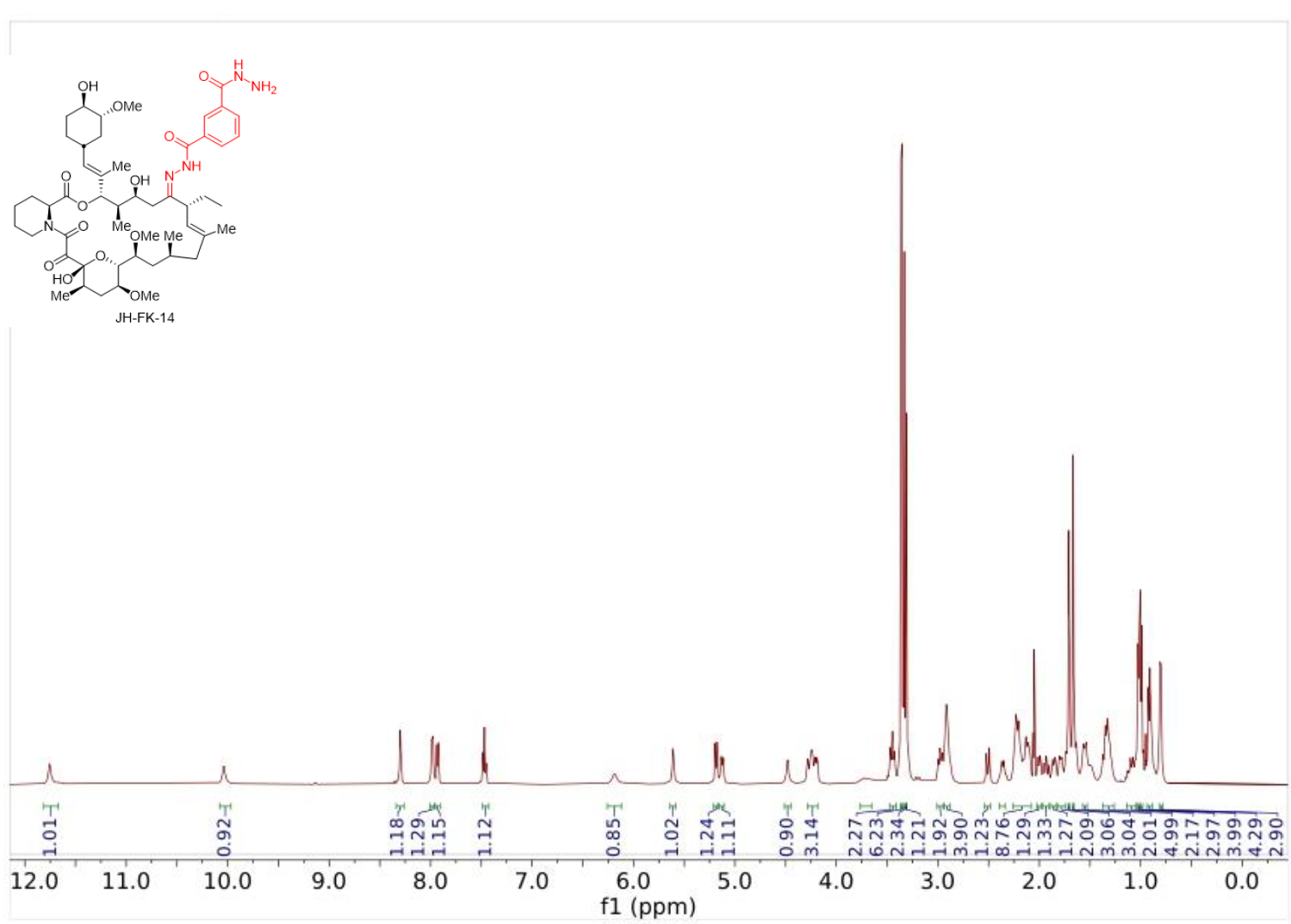

**JH-FK-15**: JH-FK-15 was synthesized from FK520 (100 mg, 0.13 mmol) and 2-naphthoic acid hydrazide (6 eq, 141 mg, 0.76 mmol) in MeOH (1.3 mL, 0.1 M) according to General Procedure B. Purification by column chromatography (silica gel, CH_2_Cl_2_/MeOH, 25/1) afforded JH-FK-15 (57 mg, 47%) as a white solid: ^1^H NMR (500 MHz, acetone-*d_6_*): δ 11.56 (s, 1H), 8.52 (s, 1H), 8.06 (d, *J* = 9.5 Hz, 1H), 7.98 (dd, *J* = 9.0, 1.5 Hz, 1H), 7.94–7.90 (m, 2H), 7.57 (td, *J* = 2.0, 7.0 Hz, 1H), 7.54 (td, *J* = 2.0 Hz, 7.0 Hz, 1H), 5.74 (s, 2H), 5.19 (d, *J* = 9.0 Hz, 1H), 5.17 (d, *J* = 11 Hz, 1H), 4.45 (s, 1H), 4.37 (s, 1H), 4.29 (d, *J* = 14 Hz, 1H), 4.19 (d, *J* = 11 Hz, 1H), 3.67 (s, 1H), 3.49 (d, *J* = 11 Hz, 1H), 3.46 (d, *J* = 9.5 Hz, 1H), 3.36 (d, *J* = 4.0 Hz, 6H), 3.34 (s, 2H), 3.32 (s, 3H), 3.17 (br s, 1H), 3.02–2.94 (m, 2H), 2.89 (s, 1H), 2.53 (d, *J* = 14 Hz, 1H), 2.43–2.35 (m, 1H), 2.29–2.16 (m, 6H), 2.13–2.08 (m, 1H), 2.03–1.98 (m, 1H), 1.93 (t, *J* = 11 Hz, 1H), 1.90–1.79 (m, 3H), 1.77–1.73 (m, 2H), 1.71 (s, 3H), 1.69 (s, 3H), 1.65–1.47 (m, 5H), 1.39–1.27 (m, 4H), 1.05–1.02 (m, 4H), 0.96 (q, *J* = 12 Hz, 2H), 0.91–0.87 (m, 3H), 0.79 (d, *J* = 6,5 Hz, 3H).

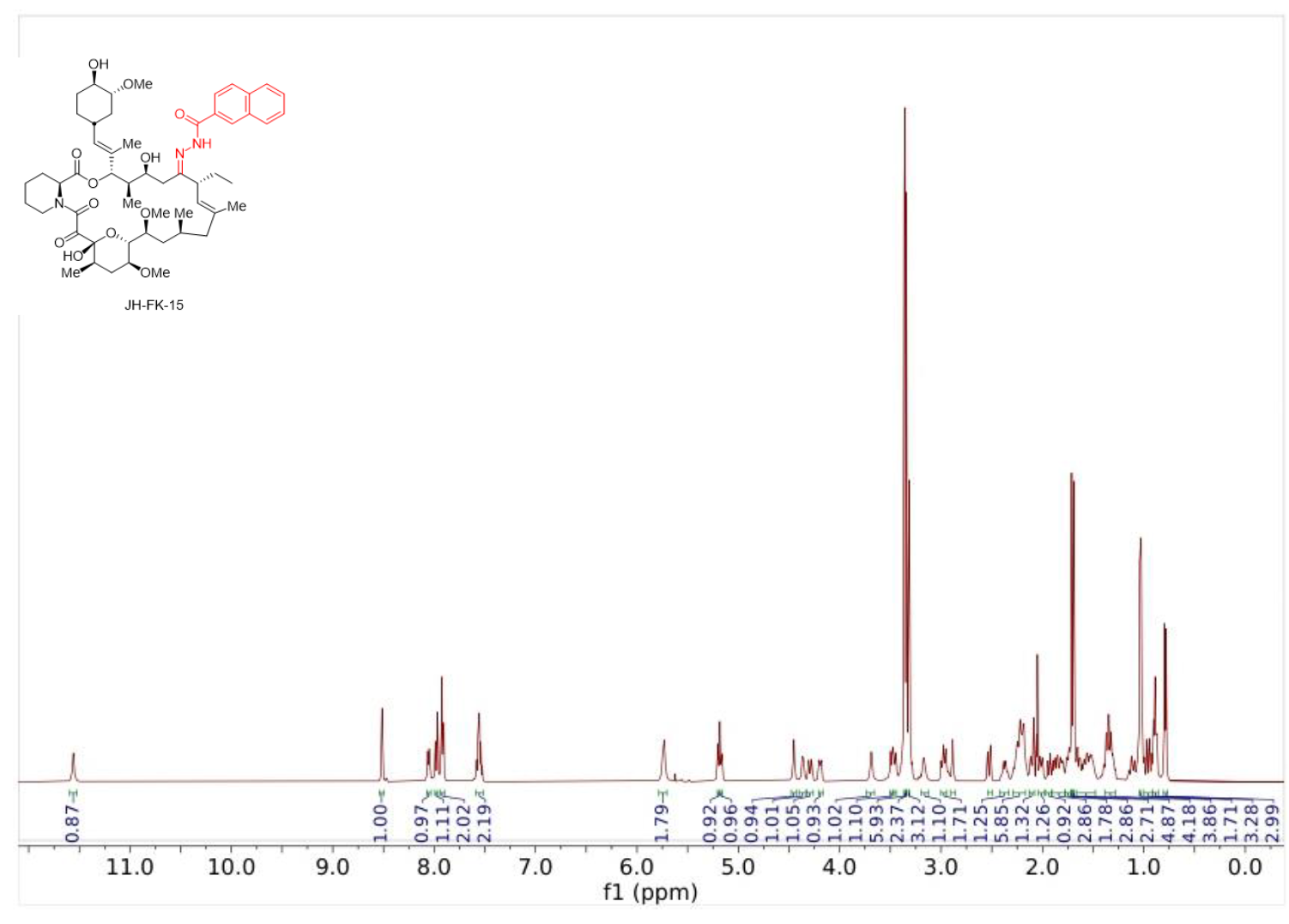

**JH-FK-16**: JH-FK-16 was synthesized from FK520 (100 mg, 0.13 mmol) and ethylhydrazine hydrochloride (6 eq, 73 mg, 0.76 mmol) in MeOH (1.3 mL, 0.1 M) according to General Procedure B. Purification by column chromatography (silica gel, CH_2_Cl_2_/MeOH, 25/1) afforded JH-FK-16 (44 mg, 42%) as a white solid (44 mg, 42%) ^1^H NMR (500 MHz, acetone-*d_6_*): δ 7.62 (s, 1H), 6.14 (t, *J* = 6.0 Hz, 1H), 5.33 (s, 1H), 4.99 (d, *J* = 8.0 Hz, 1H), 4.74–4.68 (m, 2H), 4.41 (d, *J* = 6.0 Hz, 1H), 4.30 (d, *J* = 15 Hz, 1H), 3.90 (d, *J* = 7.5 Hz, 1H), 3.61 (d, *J* = 3.0 Hz, 1H), 3.60 (dd, *J* = 3.5, 7.5 Hz, 1H), 3.45 (d, *J* = 3.5 Hz, 1H), 3.44 (s, 1H), 3.36–3.35 (m, 7H), 3.34–3.28 (m, 4H), 3.12–3.06 (m, 2H), 3.00–2.94 (m, 4H), 2.41–2.33 (m, 2H), 2.27–2.14 (m, 5H), 2.11 (s, 1H), 1.99–1.94 (m, 1H), 1.90–1.77 (m, 6H), 1.75 (s, 3H), 1.66 (s, 3H), 1.61 (s, 1H) 1.59–1.546 (m, 2H), 1.40–1.29 (m, 9H), 1.14 (t, *J* = 7.0 Hz, 3H), 1.03 (d, *J* = 6.0 Hz, 2H), 0.92 (s, 1H), 0.91 (s, 1H), 0.87–0.85 (m, 1H), 0.84 (d, *J* = 3.0 Hz, 2H), 0.83–0.81 (m, 2H).

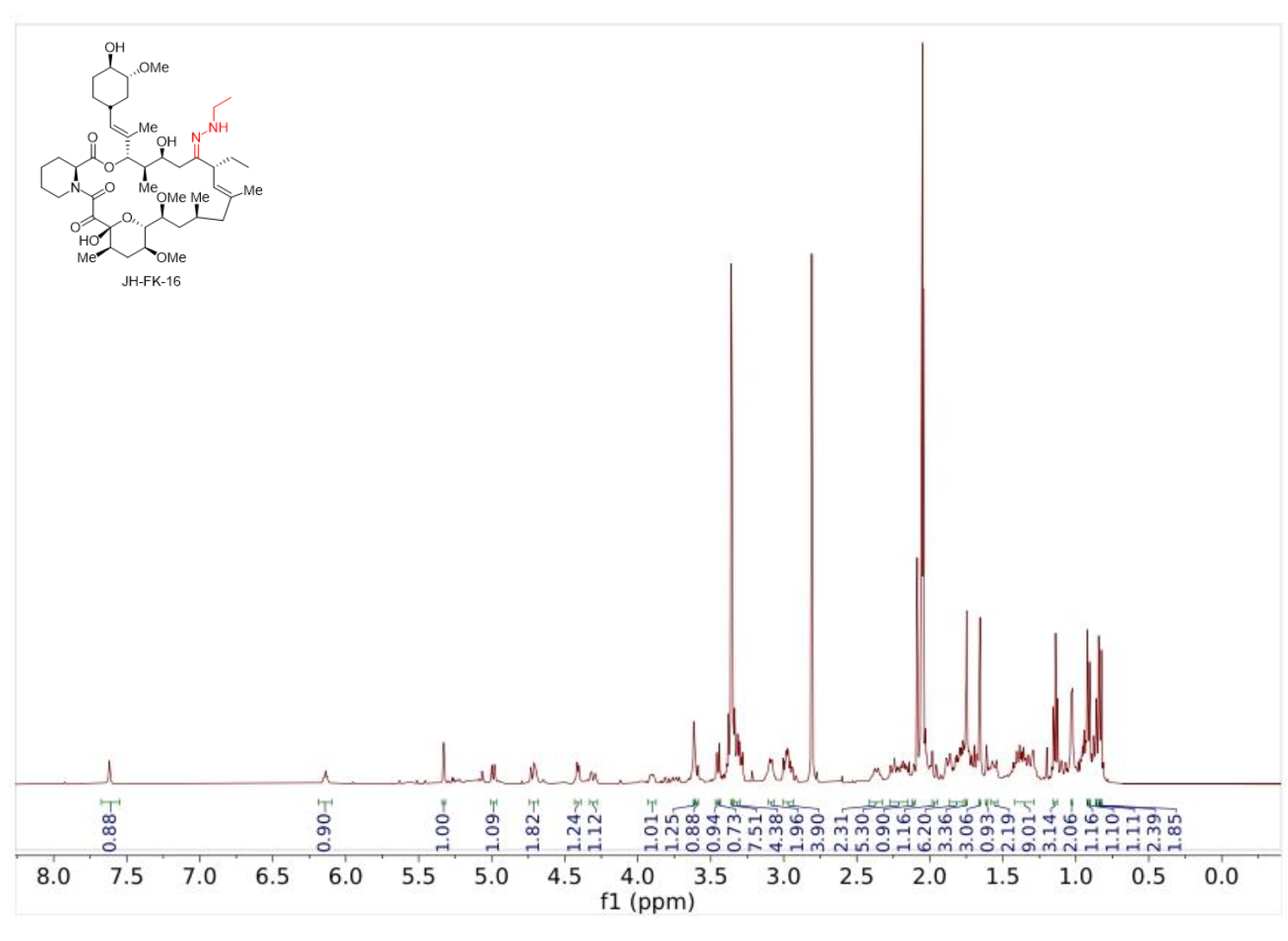

**JH-FK-17**: JH-FK-17 was synthesized from FK520 (100 mg, 0.13 mmol) and 4-fluorobenzoic acid hydrazide (6 eq, 147 mg, 0.76 mmol) in MeOH (1.3 mL, 0.1 M) according to General Procedure B. Purification by column chromatography (silica gel, CH_2_Cl_2_/MeOH, 25/1) afforded JH-FK-17 (53 mg, 45%) as a white solid: ^1^H NMR (500 MHz, acetone-*d_6_*): δ 11.29 (s, 1H), 7.83 (dd, *J* = 5.5, 9.0 Hz, 2H), 7.05 (d, *J* = 10 Hz, 1H), 7.03 (d, *J* = 9 Hz, 1H), 5.49 (s, 1H), 5.05 (d, *J* = 11 Hz, 1H), 5.02 (d, *J* = 9.0 Hz, 1H), 4.36 (s, 1H), 4.19–4.14 (m, 2H), 4.02 (d, *J* = 6.0 Hz, 1H), 3.63 (s, 1H), 3.36–3.29 (m, 2H), 3.25–3.23 (m, 4H), 3.21 (s, 2H), 3.19–3.18 (m, 10H), 3.11 (br s, 3H), 2.37 (d, *J* = 14 Hz, 1H), 2.29–2.21 (m, 1H), 2.10–1.99 (m, 6H), 1.90–1.86 (m, 1H), 1.81–1.59 (m, 5H), 1.56 (s, 3H), 1.55 (s, 3H), 1.53–1.41 (m, 4H), 1.26–1.16 (m, 4H), 1.02–0.94 (m, 1H), 0.89 (d, *J* = 5.5 Hz, 3H), 0.88 (d, *J* = 7.5 Hz, 2H), 0.84–0.81 (m, 1H), 0.77 (t, *J* = 7.5 Hz, 3H), 0.67 (d, *J* = 6.5 Hz, 2H).

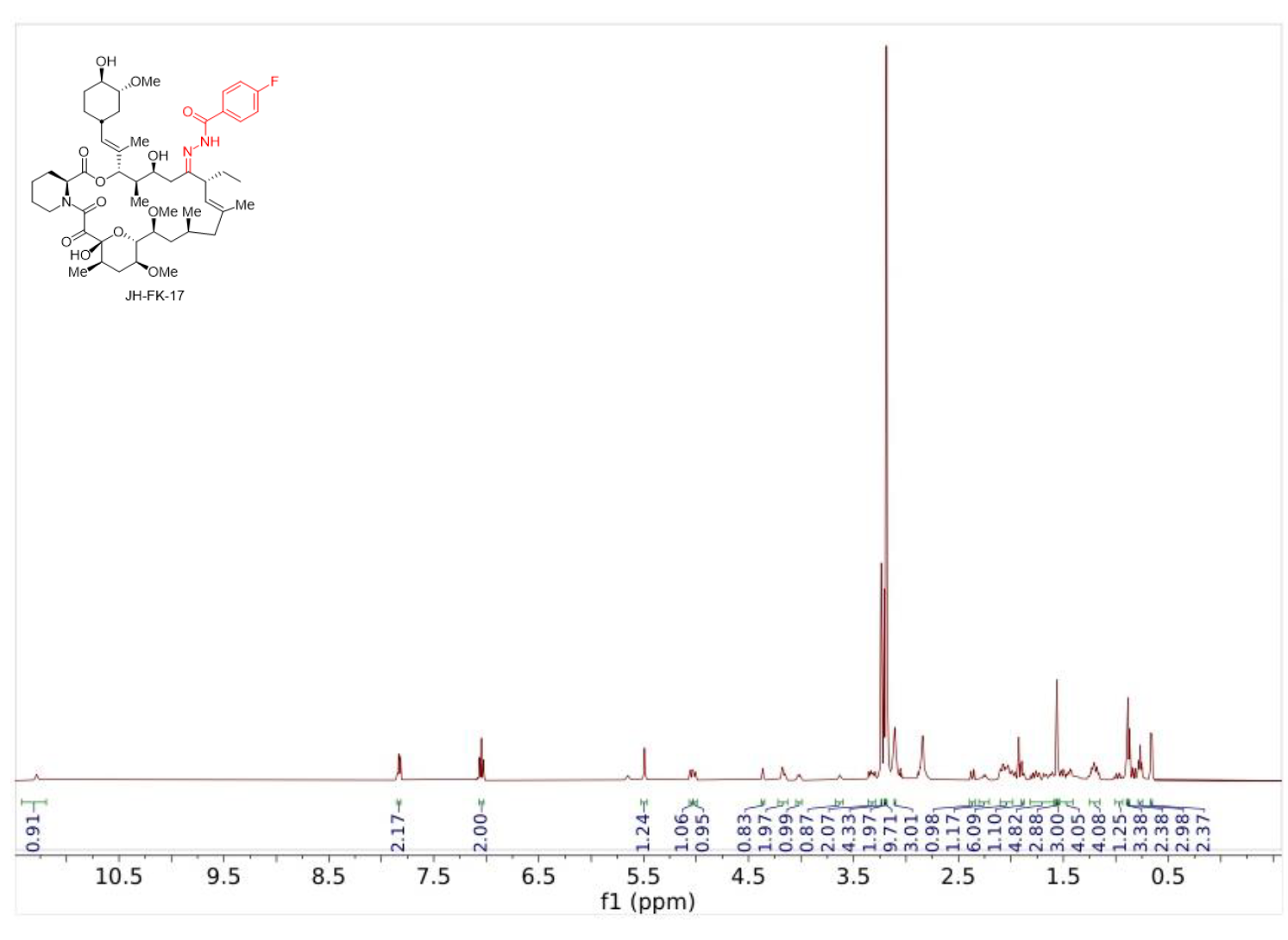

**JH-FK-20**: JH-FK-20 was synthesized from FK520 (100 mg, 0.13 mmol) and 4-aminobenzoic hydrazide (6 eq, 114 mg, 0.76 mmol) in MeOH (1.3 mL, 0.1 M) according to General Procedure B. Purification by column chromatography (silica gel, CH_2_Cl_2_/MeOH, 25/1) afforded JH-FK-20 (50 mg, 43%) as a white solid (50 mg, 43%): ^1^H NMR (500 MHz, acetone-*d_6_*): δ 11.01 (s, 1H), 7.68 (d, *J* = 9.5 Hz, 2H), 6.62 (d, *J* = 9.0 Hz, 2H), 5.54 (s, 1H), 5.40 (s, 1H), 5.18 (d, *J* = 9.5 Hz, 1H), 5.15 (d, *J* = 9.5 Hz, 1H), 5.06 (s, 2H), 4.56 (s, 1H), 4.32–4.27 (m, 2H), 4.14–4.08 (m, 1H), 3.63 (d, *J* = 2.5 Hz, 1H), 3.47 (d, *J* = 9.0 Hz, 1H), 3.45 (d, *J* = 7.5 Hz, 1H), 3.37–3.35 (m, 6H), 3.33 (s, 3H), 3.32–3.29 (m, 2H) 3.00–2.94 (m, 2H), 2.45 (d, *J* = 14 Hz, 1H), 2.41–2.34 (m, 1H), 2.25–2.08 (m, 8H), 2.02–1.98 (m, 1H), 1.95–1.73 (m, 6H), 1.69 (s, 3H), 1.68 (s, 3H), 1.66–1.50 (m, 5H), 1.39–1.29 (m, 4H), 1.14–1.05 (m, 2H), 1.01 (d, *J* = 2.5 Hz, 2H), 1.00 (d, *J* = 3.0 Hz, 2H), 0.95 (q, *J* = 13 Hz, 1H), 0.89 (t, *J* = 7.0 Hz, 3H), 0.79 (d, *J* = 6.5 Hz, 3H).

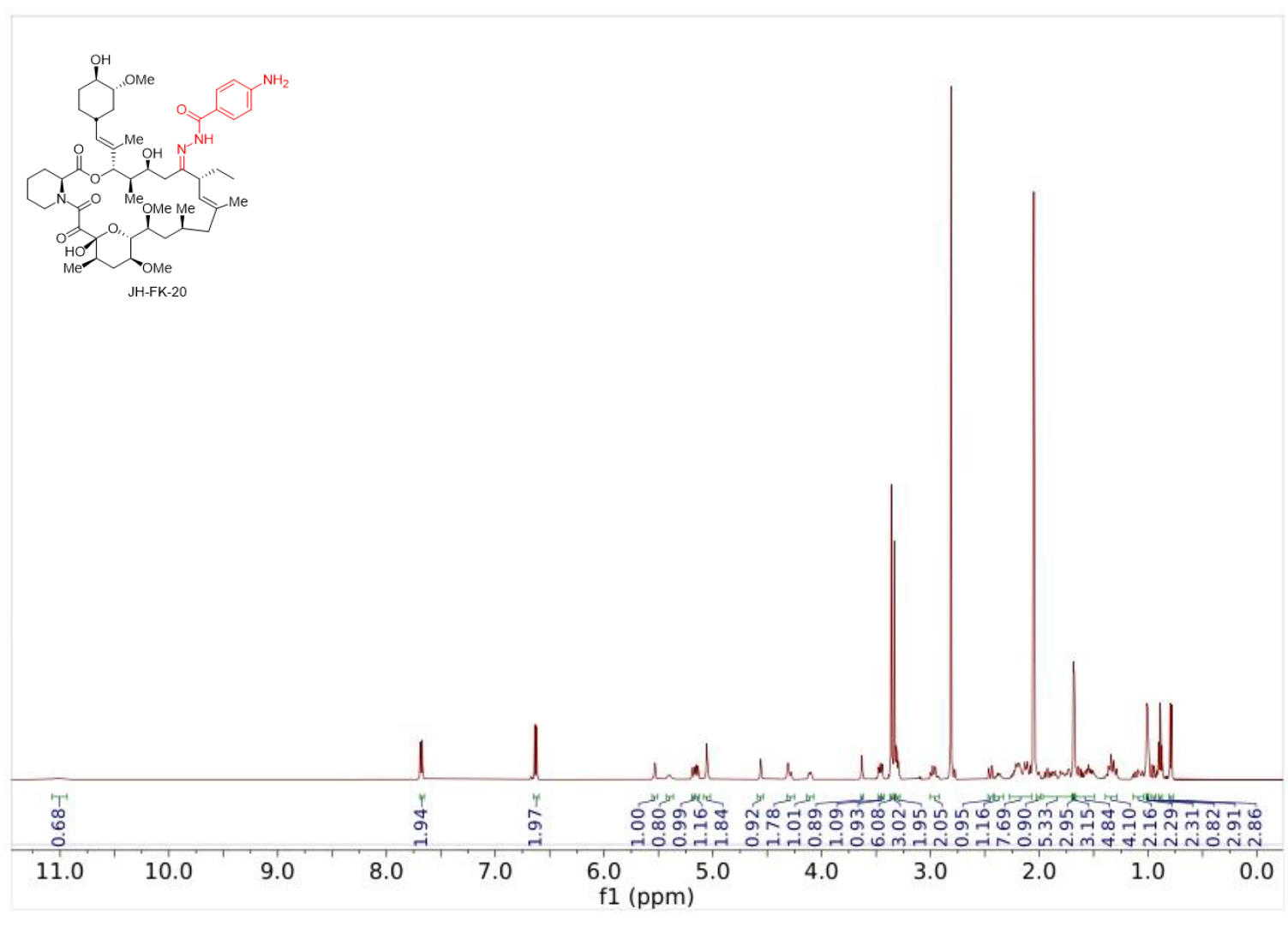

**JH-FK-21**: JH-FK-21 was synthesized from FK520 (100 mg, 0.13 mmol) and carbohydrazide (6 eq, 68.1 mg, 0.76 mmol) in MeOH (1.3 mL, 0.1 M) according to General Procedure B. Purification by column chromatography (silica gel, CH_2_Cl_2_/MeOH, 25/1) afforded JH-FK-21 (37 mg, 34%) as a white solid (37 mg, 34%): ^1^H NMR (500 MHz, acetone-*d_6_*): δ 10.88 (m, 1H), 9.23 (m, 1H), 5.24 (m, 2H), 5.12 (d, *J* = 9.0 Hz, 2H), 5.01–4.57 (m, 2H), 4.33 (d, *J* = 18 Hz, 1H), 3.96–3.80 (m, 1H), 3.75–3.70 (m, 1H), 3.58 (s, 1H), 3.48–3.41 (m, 1H), 3.37 (s, 3H), 3.34 (s, 2H), 3.33 (s, 3H), 3.31 (s, 3H), 3.30 (s, 3H), 3.26–3.21 (m, 3H), 2.95 (s, 4H), 2.45–2.35 (m, 1H), 2.28–2.15 (m, 5H), 2.00–1.93 (m, 5H), 1.91–1.89 (m, 3H), 1.77–1.71 (m, 3H), 1.70 (s, 3H), 1.61–1.48 (m, 5H), 1.41–1.28 (m, 3H), 0.99 (d, *J* = 7.0 Hz, 3H), 0.96–0.84 (m, 8H).

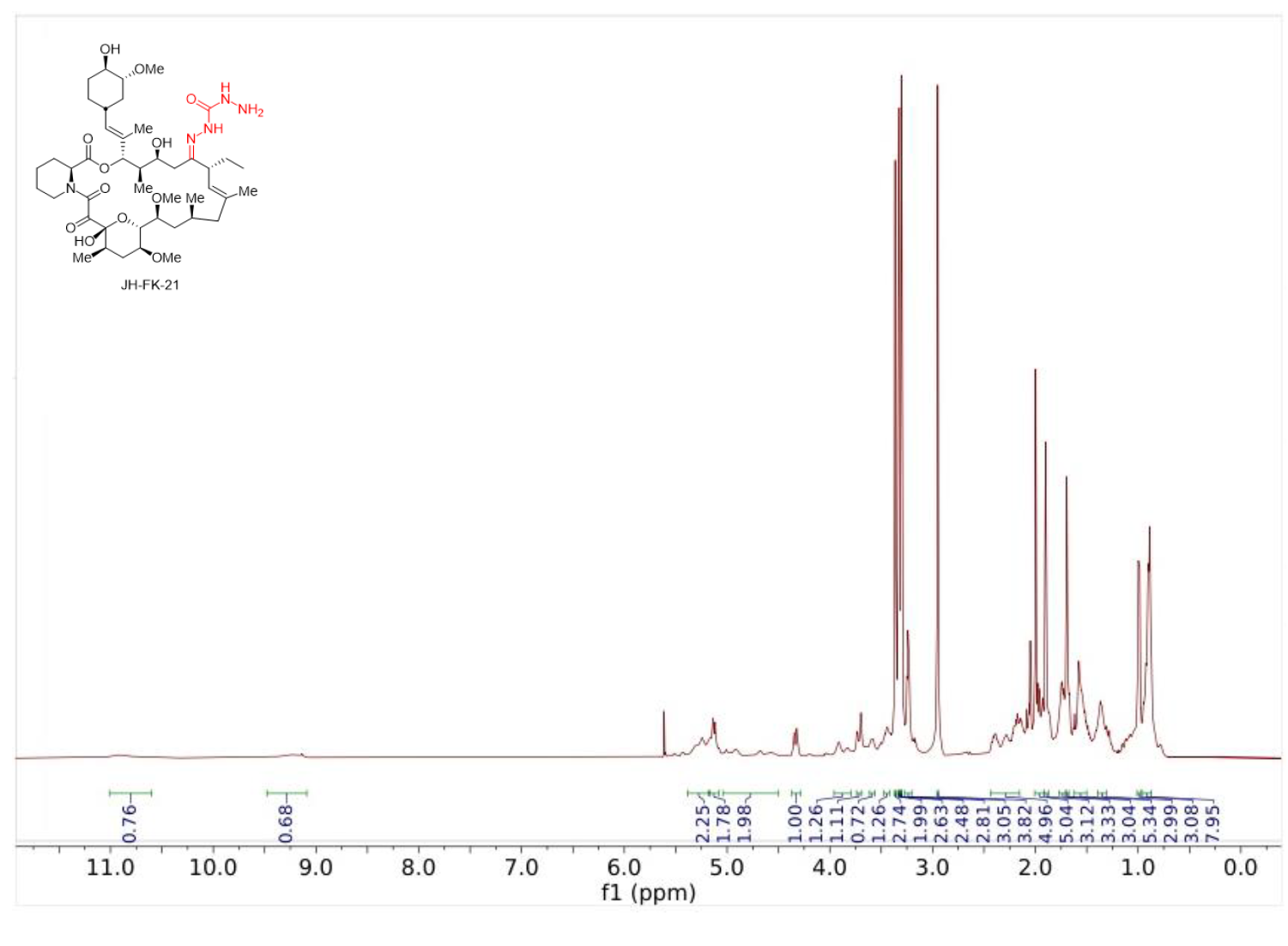

**JH-FK-22**: JH-FK-22 was synthesized from FK520 (100 mg, 0.13 mmol) and 2-hydroxyacetic acid hydrazide (6 eq, 45 mg, 0.76 mmol) in MeOH (1.3 mL, 0.1 M) according to General Procedure B. Purification by column chromatography (silica gel, CH_2_Cl_2_/MeOH, 25/1) afforded JH-FK-22 (48 mg, 41%) as a white solid: ^1^H NMR (500 MHz, acetone-*d_6_*): δ 11.01 (s, 1H), 10.27 (s, 1H), 5.53–5.45 (s, 1H), 5.20–5.15 (m, 2H), 5.02 (s, 1H), 4.46 (s, 1H), 4.41 (d, *J* = 5.0 Hz, 1H), 4.36 (s, 1H), 4.34 (s, 1H), 4.32–4.26 (m, 1H), 4.21–4.15 (m, 1H), 4.09–4.02 (m, 1H), 3.65 (s, 1H), 3.56 (d, *J* = 10 Hz, 1H), 3.48 (d, *J* = 8.5 Hz, 1H), 3.45–3.41 (m, 3H), 3.39–3.37 (m, 6H), 3.35 (s, 3H), 3.33–3.32 (m, 1H), 3.21 (q, *J* = 7.5 Hz, 1H), 3.11 (m, 1H), 2.99 (m, 1H), 2.38 (m, 2H), 2.26–2.10 (m, 6H), 1.97 (d, *J* = 2.0 Hz, 1H), 1.92–1.86 (m, 2H), 1.82–1.76 (m, 2H), 1.69–1.67 (m, 4H), 1.66 (s, 1H), 1.64–1.57 (m, 4H), 1.39–1.29 (m, 4H), 1.19–1.08 (m, 2H), 1.04–0.97 (m, 6H), 0.91–0.84 (m, 7H).

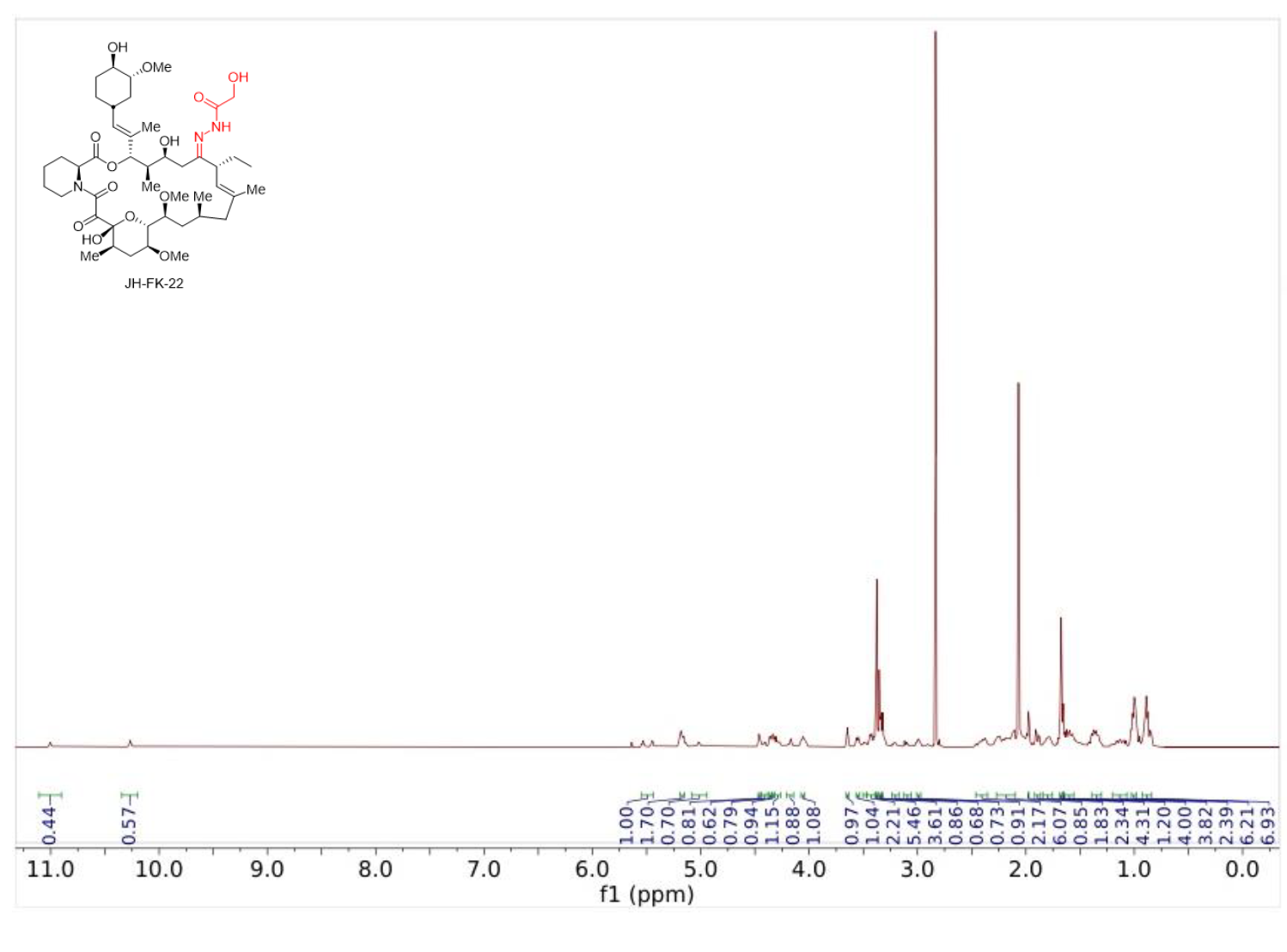

**JH-FK-23**: JH-FK-23 was synthesized from FK520 (100 mg, 0.13 mmol) and methanesulfonyl hydrazide (6 eq, 83.3 mg, 0.76 mmol) in MeOH (1.3 mL, 0.1 M) according to General Procedure B. Purification by column chromatography (silica gel, CH_2_Cl_2_/MeOH, 25/1) afforded JH-FK-23 (58 mg, 52%) as a white solid: ^1^H NMR (500 MHz, acetone-*d_6_*): δ 9.51 (s, 1H), 5.40 (d, *J* = 3.5 Hz, 1H), 5.30 (s, 1H), 5.14 (t, *J* = 7.5 Hz, 2H), 4.68 (s, 1H), 4.38 (d, *J* = 4.0 Hz, 1H), 4.33 (d, *J* = 12 Hz, 1H), 4.06 (d, *J* = 9.5 Hz, 1H), 3.65 (d, *J* = 3.0 Hz, 1H), 3.49 (d, *J* = 9.5 Hz, 1H), 3.46 (d, *J* = 9.5 Hz, 1H), 3.40–3.37 (m, 1H), 3.37 (s, 2H), 3.36 (s, 3H), 3.34 (s, 3H), 3.32–3.30 (m, 1H), 3.22 (q, *J*= 8.0 Hz, 1H), 3.03–2.99 (m, 1H), 2.98 (s, 3H), 2.96–2.93 (m, 1H), 2.40–2.34 (m, 2H), 2.29–2.23 (m, 1H), 2.20–2.10 (m, 5H), 2.03–1.98 (m, 1H), 1.95–1.91 (m, 1H), 1.90–1.84 (m, 2H), 1.81 (t, *J* = 13 Hz, 2H), 1.73–1.70 (m, 1H), 1.67 (s, 3H), 1.64 (s, 3H), 1.61–1.51 (m, 4H), 1.38–1.28 (m, 4H), 1.10–1.03 (m, 3H), 0.99 (d, *J* = 6.5 Hz, 3H), 0.98 (d, *J* = 8.0 Hz, 3H), 0.96–0.93 (m, 1H), 0.89 (t, *J* = 7.5 Hz, 4H), 0.86 (d, *J* = 6.5 Hz, 2H).

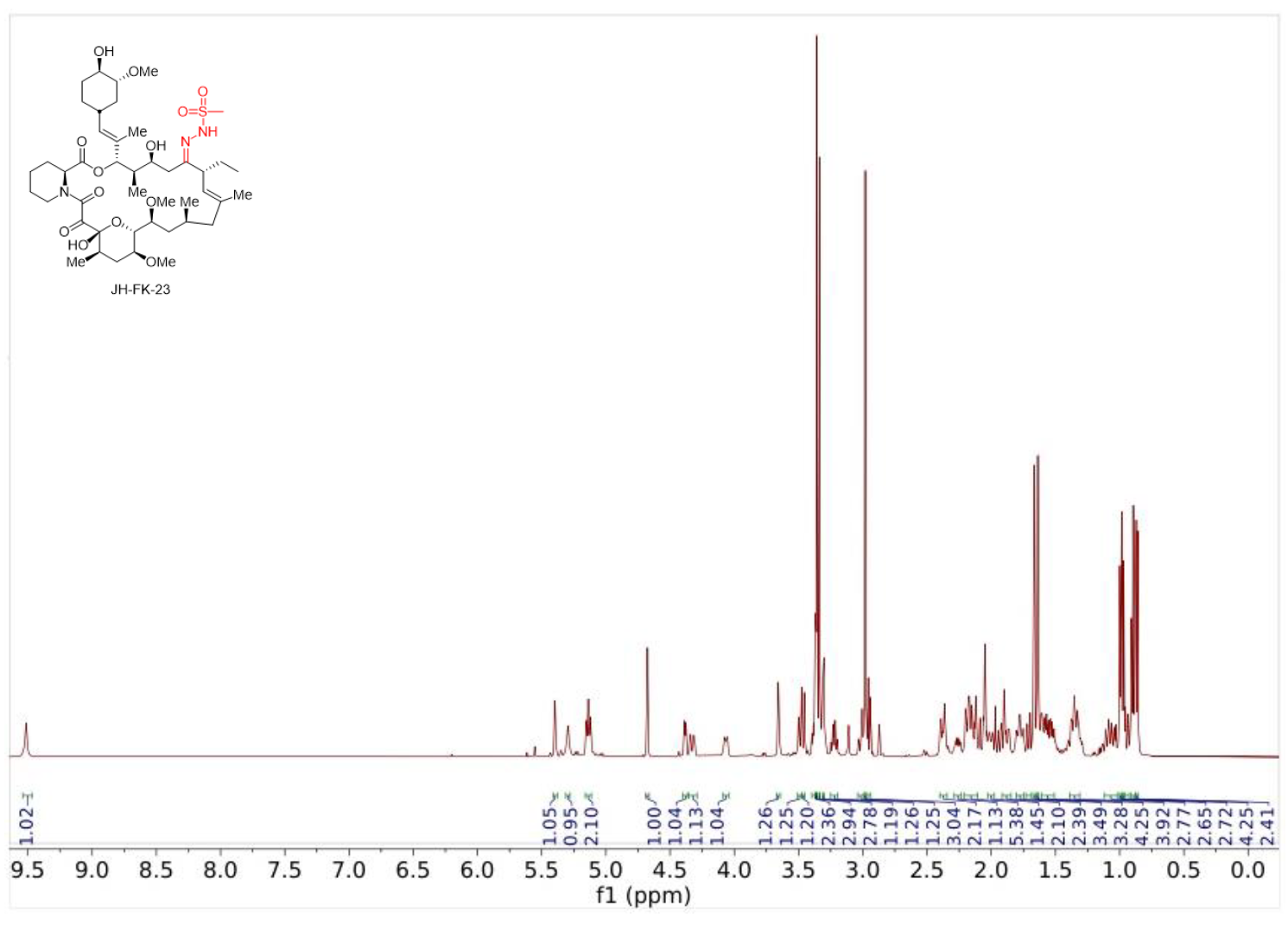

Table S1. High-resolution mass spectrometry (HRMS) data for FK520 analogs

| Analog | Elemental Composition | Calculated Mass | Observed Mass | Error (ppm) | Ion |
| --- | --- | --- | --- | --- | --- |
| JH-FK-08 | C_45_H_73_N_3_O_13_ | 864.5222 | 864.5217 | -0.6 | [M+H]^+^ |
| JH-FK-09 | C_49_H_74_N_4_O_12_ | 911.5376 | 911.5378 | 0.2 | [M+H]^+^ |
| JH-FK-12 | C_44_H_72_N_4_O_12_ | 849.5220 | 849.5223 | 0.4 | [M+H]^+^ |
| JF-FK-13 | C_50_H_77_N_3_O_13_S | 960.5250 | 960.5261 | 0.1 | [M+H]^+^ |
| JH-FK-14 | C_51_H_77_N_5_O_13_ | 968.5591 | 968.5575 | -1.7 | [M+H]^+^ |
| JH-FK-15 | C_54_H_77_N_3_O_12_ | 960.5580 | 960.5579 | -0.1 | [M+H]^+^ |
| JH-FK-16 | C_45_H_75_N_3_O_11_ | 834.5474 | 834.5482 | 1.0 | [M+H]^+^ |
| JH-FK-17 | C_50_H_74_FN_3_O_12_ | 928.5329 | 928.5341 | 1.3 | [M+H]^+^ |
| JH-FK-20 | C_50_H_76_N_4_O_12_ | 925.5533 | 925.5537 | 0.4 | [M+H]^+^ |
| JH-FK-21 | C_44_H_73_N_5_O_12_ | 864.5329 | 864.5319 | -1.2 | [M+H]^+^ |
| JH-FK-22 | C_45_H_73_N_3_O_13_ | 864.5216 | 864.5214 | -0.2 | [M+H]^+^ |
| JH-FK-23 | C_44_H_73_N_3_O_13_S | 884.4937 | 884.4930 | -0.8 | [M+H]^+^ |

A

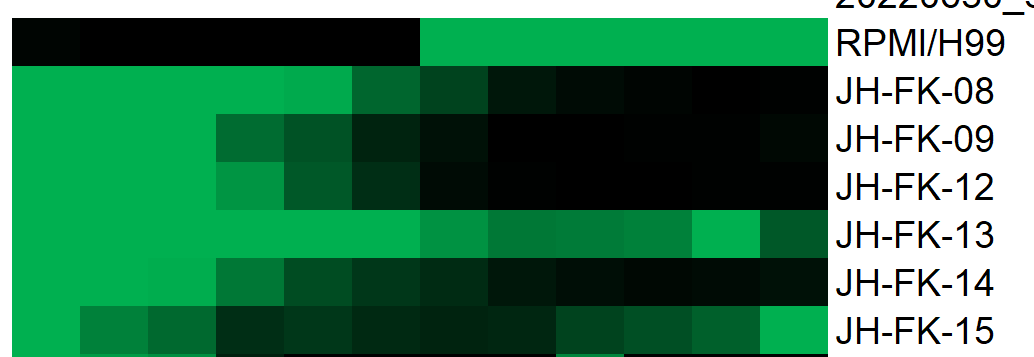

B

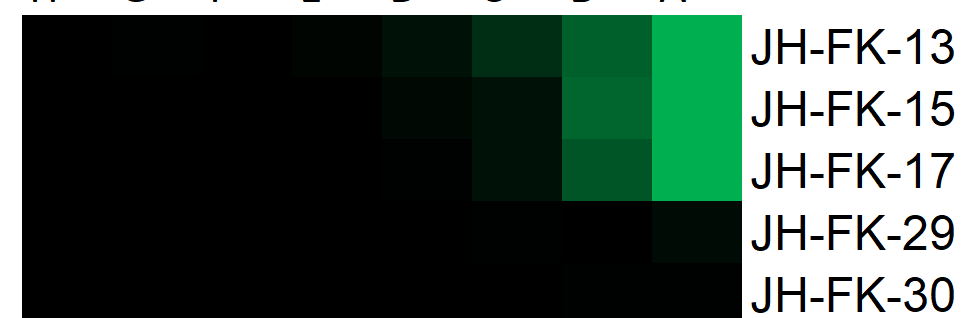

**Fig. S1.** Low-solubility compounds in MIC assay. Representative image of MIC assay read by plate reader with growth indicated on a range from 0.04 to 0.1 OD, black to green coloration. Starting concentration 50 μg/mL for all drugs. (*A*) H99 cultured in wells, and (*B*) wells contain RPMI media and no fungal cells with green coloration indicative of false positive growth caused by drug insolubility.

| **Table S2.** Antifungal activity of JH-FK-08 against natural and mutant strains | | |
| --- | --- | --- |
| **Fungal strain** | **Background** | **MIC / MEC (μg/mL)** |
| KN99α | *Cryptococcus neoformans* | <0.4 |
| *frr1*Δ | KN99α *frr1*Δ::*NAT* | >25 |
| JEC21 | *Cryptococcus deneoformans* | <0.4 |
| C21F2 | JEC21 *CNB1-1* | >25 |
| C21F3 | JEC21 *frr1* | >25 |
| SC5314 | *Candida albicans* | <0.4 |
| YAG171 | SC5314 *rbp1*Δ::*URA3/rbp1*Δ::*HISG* | >25 |
| YAG237 | SC5314 *CNB1-1/CNB1* | >25 |
| *akuB*^Ku80^ | *Aspergillus fumigatus* | 1.25 |
| Δ*fkbp12* | akuB^Ku80^ *fkbp12*Δ | >20 |
| H99 | *Cryptococcus neoformans* | >4 |
| MIC_90_ was collected at 37°C for KN99 and related strains. MIC_90_ was collected at 30°C for H99 JEC21 strains grown for 72 hours with 5 times starting amount of cells. MIC_80_ was collected at 30°C with 2 μg/mL supplementary fluconazole grown for 72 hours for *C. albicans.* MEC was collected for *A. fumigatus* strains. | | |

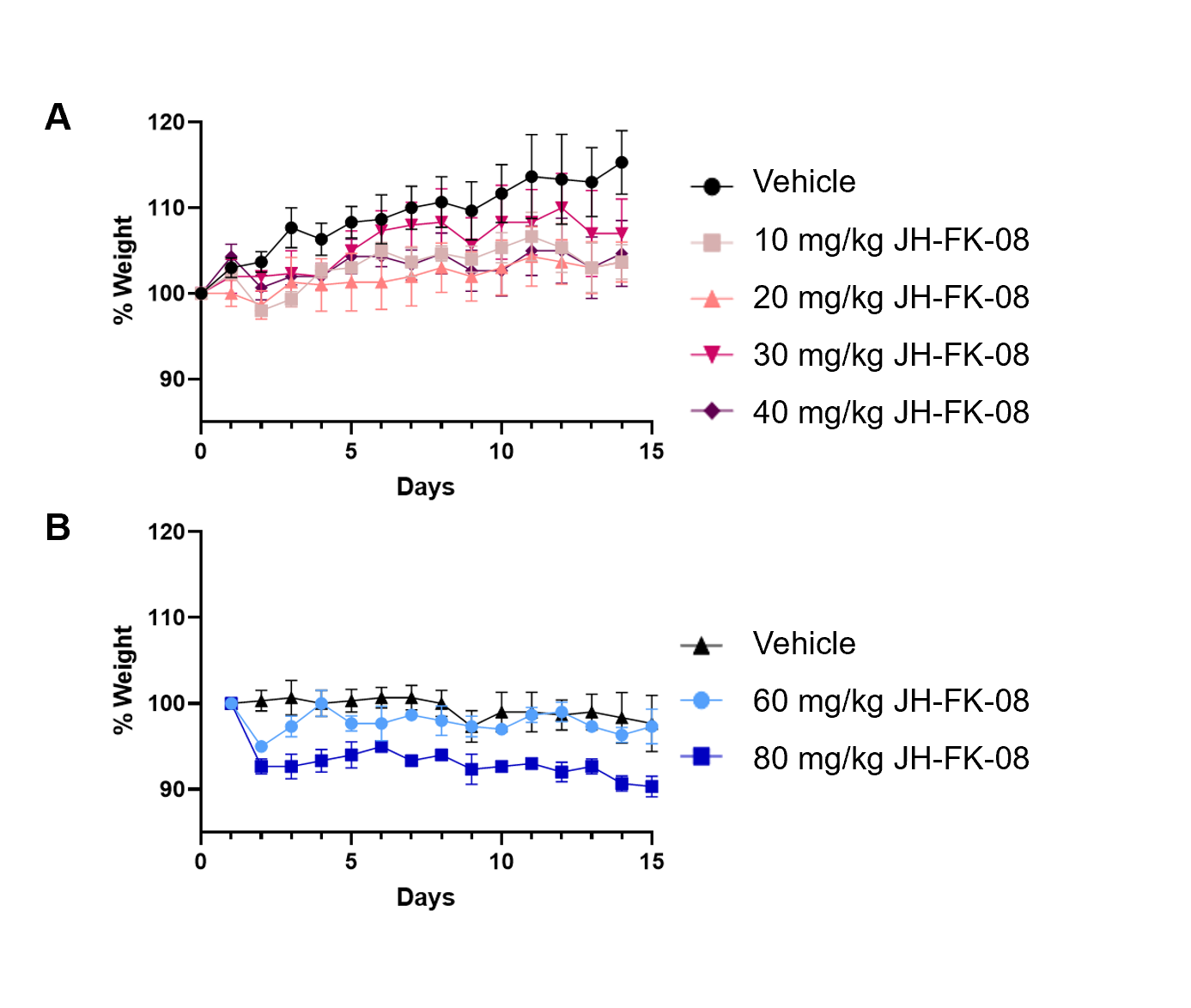

**Fig S2.** *In vivo* tolerability of JH-FK-08. Maximum tolerated dose study of female A/J mice (n=3). Animals received once daily (*A*) and twice daily (*B*) dosing, and weight over time was plotted as percent change from initial body weight.

**Table S3.** Pharmacokinetic analysis of FK506 and JH-FK-08.

|  | **FK506 Females** | | **FK506 Males** | | **JH-FK-08 Females** | | **JH-FK-08 Males** | |
| --- | --- | --- | --- | --- | --- | --- | --- | --- |
|  | Value | st dev | Value | st dev | Value | st dev | Value | st dev |
| **T_max_** [h] | 1.11 | 0.67 | 0.75 | 0.00 | 2.50 | 0.87 | 1.25 | 0.43 |
| **C_max_** [μg/mL] | 5.96 | 1.11 | 4.51 | 0.62 | 7.50 | 1.33 | 4.07 | 1.22 |
| **AUC** (area under the curve) [μg mL-1 h] | 27.10 | 7.28 | 15.17 | 5.72 | 31.42 | 4.32 | 15.64 | 5.05 |
| **t_1/2_ (3-12h)** (half-life) [h] | 1.35 | 0.16 | 1.48 | 0.14 | 1.14 | 0.02 | 1.29 | 0.05 |
| **t_1/2_ (12-24h)** (half-life) [h] | 3.59 | 0.95 | 2.46 | 0.22 | 1.88 | 0.03 | 2.26 | 0.29 |
| **Cl/F** (clearance as Dose/AUCINF) [mL h^-1^ kg] | 1555 | 452 | 2961 | 1310 | 1291 | 192 | 2786 | 1065 |

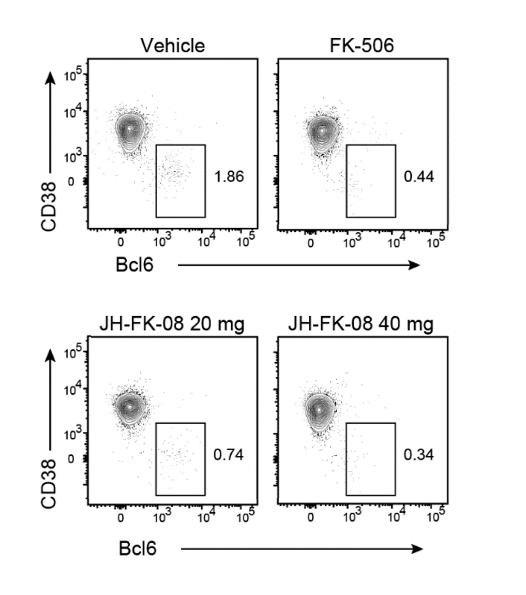

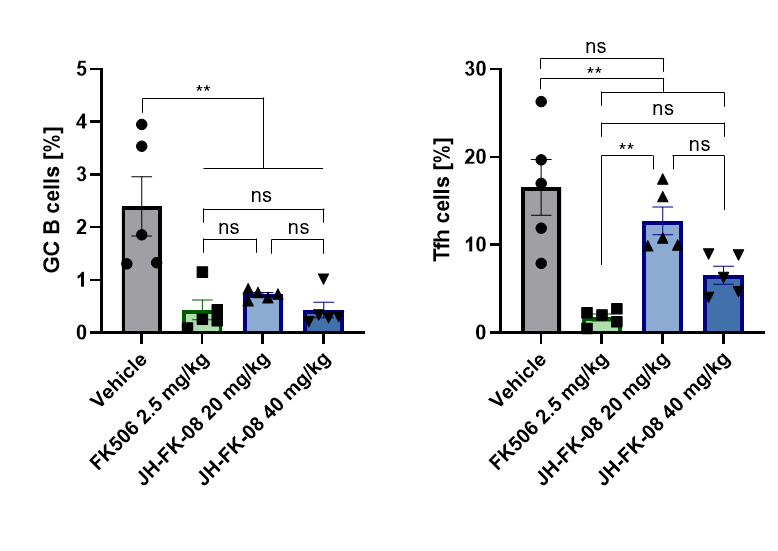

**A**

**B**

**Fig. S3.** *In vivo* immunosuppression of JH-FK-08 compared to FK506. (*A*) Shown here is the production of germinal center B (BC) cells during immune response in C57BL/6 female mice. (*B*) Shown here is a representative flow cytometry plots showing gating to indicate region containing BC cells among CD19^+^ B cells. Mice received twice daily treatment of vehicle, FK506 (2.5 mg/kg), JH-FK-08 (20 mg/kg), or JH-FK-08 (40 mg/kg) (n=5; **, p<0.001, ordinary One-way ANOVA with Tukey’s test).

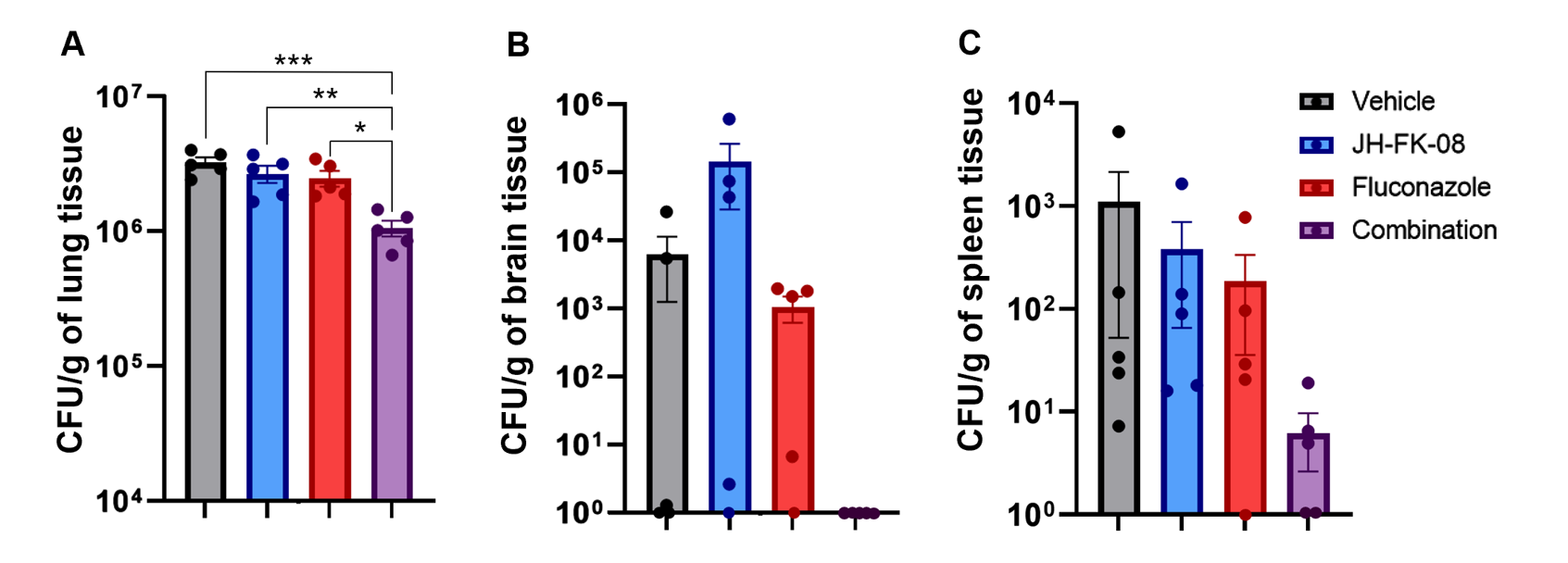

**Fig. S4.** JH-FK-08 and fluconazole treatment reduced organ fungal burden in infected animals. Female A/J mice (n=5) were infected with *C. neoformans* strain H99 through intranasal inoculation and treated daily with vehicle, 12 mg/kg fluconazole, 40 mg/kg JH-FK-08, or combination through i.p. dosing. Fungal burden levels in lung (*A*), brain (*B*), and spleen (*C*) were assessed after 14 days of treatment. ( *, p=0.02; **, p=0.008; ***, p=0.0005, ordinary One-way ANOVA with Tukey’s test, error bars represent SEM).

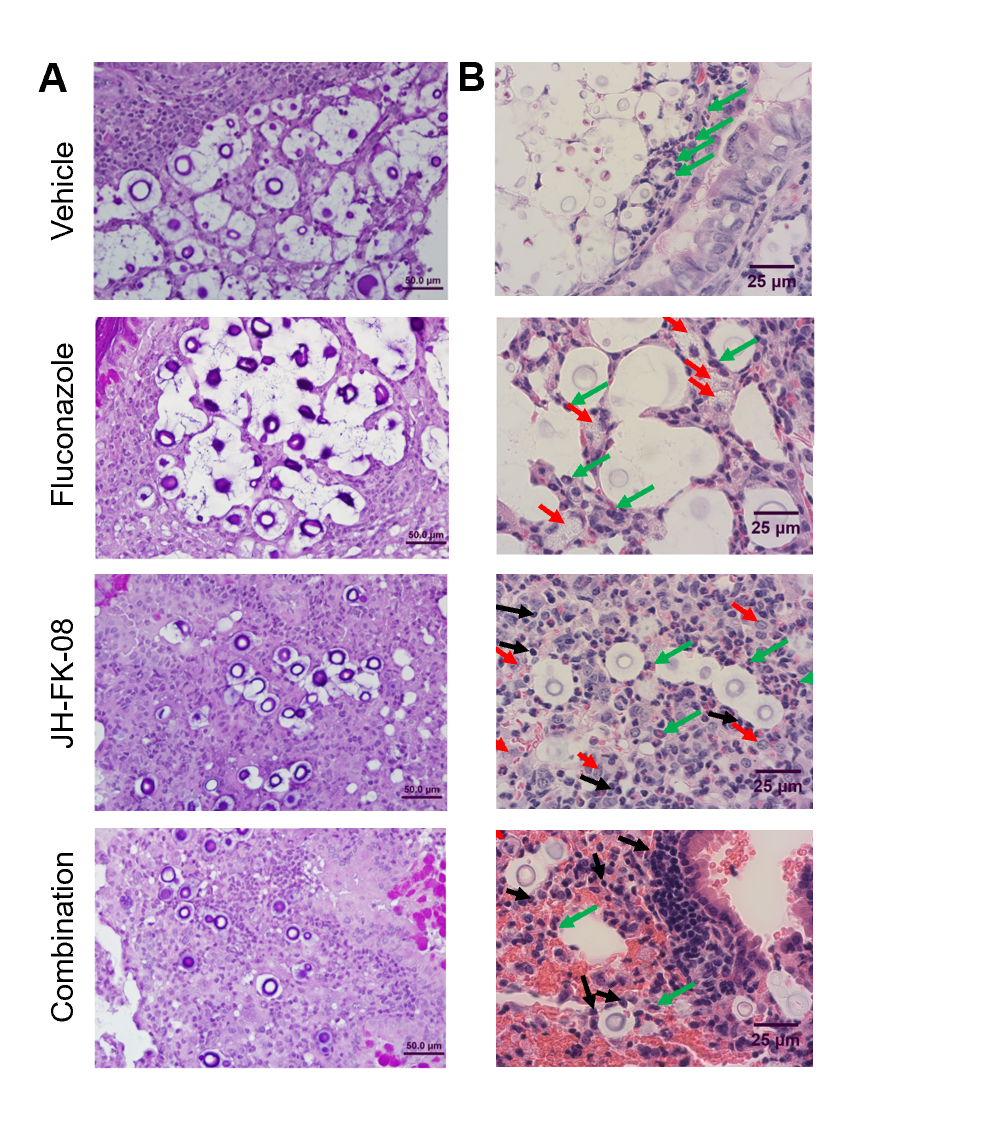

**Fig. S5.** Histopathological analysis of lung tissue from animals treated with JH-FK-08 alone or in combination with fluconazole. Representative images of lung tissue from infected airways in various treatment groups stained with (*A*) periodic acid Schiff (PAS) staining or (*B*) hematoxylin and eosin (H&E). The inflammatory response to *Cryptococcus* is similar in all treated groups and composed of a prominent granulomatous reaction, better seen at lower magnification (100X) on the left column. Detail of the immune cell types present within this inflammatory response is better seen at higher magnification (630X) in the right column. Black arrows indicate lymphocytes, red arrows macrophages, and green arrows granulocytes (n=2).
